# Spatial proteomics identifies a novel CRTC-dependent viral sensing pathway that stimulates production of Interleukin-11

**DOI:** 10.1101/2024.04.04.588067

**Authors:** Benjamin J. Ravenhill, Marisa Oliveira, George Wood, Ying Di, Colin T.R. Davies, Yongxu Lu, Robin Antrobus, Gill Elliott, Nerea Irigoyen, David J. Hughes, Paul Lyons, Betty Chung, Georg H.H Borner, Michael P. Weekes

## Abstract

Appropriate cellular recognition of viruses is essential for the generation of effective innate and adaptive antiviral immunity. Viral sensors and their signalling components thus provide a crucial first line of host defence. Many exhibit subcellular relocalisation upon activation, triggering expression of interferon and antiviral genes. To identify novel signalling factors we analysed protein relocalisation on a global scale during viral infection. CREB Regulated Transcription Coactivators-2 and 3 (CRTC2/3) exhibited early cytoplasmic-to-nuclear translocation upon a diversity of viral stimuli, in diverse cell types. This movement was depended on Mitochondrial Antiviral Signalling Protein (MAVS), cyclo-oxygenase proteins and protein kinase A. We identify a key effect of transcription stimulated by CRTC2/3 translocation as production of the pro-fibrogenic cytokine interleukin-11. This may be important clinically in viral infections associated with fibrosis, including SARS-CoV-2.

## INTRODUCTION

Subcellular relocation of cellular sensors and their signalling components forms a crucial first step in the recognition of any intracellular pathogen, and determines the outcome of infection by orchestrating effective antiviral immunity. For example, known RNA sensor RIG-I translocates upon activation to Mitochondrial Antiviral Signalling Protein (MAVS) on the outer mitochondrial membrane and mitochondria-associated-membrane (MAM) (*1*). Cyclic GMP-AMP Synthase (cGAS) and other sensors recognise cytosolic viral DNA, resulting in ER-to-Golgi transition of Stimulator of Interferon Genes (STING) (*2, 3*). Activation of these sensors leads to intermediate signalling and nuclear translocation of terminal signalling components IRF3 and NF-κB, triggering production of interferon (IFN) and synthesis of IFN-stimulated genes (ISGs), including proteins limiting viral replication (*4–6*).

IRF3 and NF-κB are critical final common mediators for viral, bacterial, fungal and parasite sensing pathways (*7, 8*). Systematic mapping of analogous, novel signalling pathways will therefore provide fundamental new insights into intrinsic immunity and transform our understanding of how pathogen recognition triggers immune gene expression. A detailed mechanistic understanding of known sensing/signalling components has led directly to (i) the development of vaccine adjuvants, (ii) insights into autoimmune disease and (iii) treatments for chronic inflammation and cancer (*9–12*).

We hypothesized that novel terminal signalling molecules would share the characteristic of subcellular redistribution. To identify these components on an global scale, we employed proteomic ‘subcellular profiling’(*13, 14*) to globally quantify subcellular protein redistribution in cells infected with the model RNA virus Sendai, which is a particularly potent RIG-I agonist due to the production of defective-interfering RNAs (*15, 16*).

## RESULTS

### Subcellular profiling identifies nucleocytoplasmic translocation of CRTC2 and CRTC3 during Sendai virus infection of human fibroblasts

Subcellular profiling (**Figure 1A**) can both identify subcellular protein location and give unbiased information about protein movement upon any stimulus (*13, 14*). We chose to infect immortalised primary human foetal foreskin fibroblasts (HFFF-TERTs) were infected as we have previously performed extensive proteomic characterization of these cells, identifying their expression of a full complement of sensing and signalling proteins, and characterising an essentially identical response to infection as their non-immortalised primary equivalents (*17*). Since overall changes in protein abundance may affect movement calculations, 8 h of infection with Sendai virus was studied, where only 87/8539 (1.0%) of proteins were induced on average >2 fold. At this time point, the nuclear translocation of our positive control IRF3 was also maximal, suggesting that sensitivity for detection of other organellar protein transitions would be optimal (**Figures S1A-D, Tables S1, S2A**).

**Figure 1.**
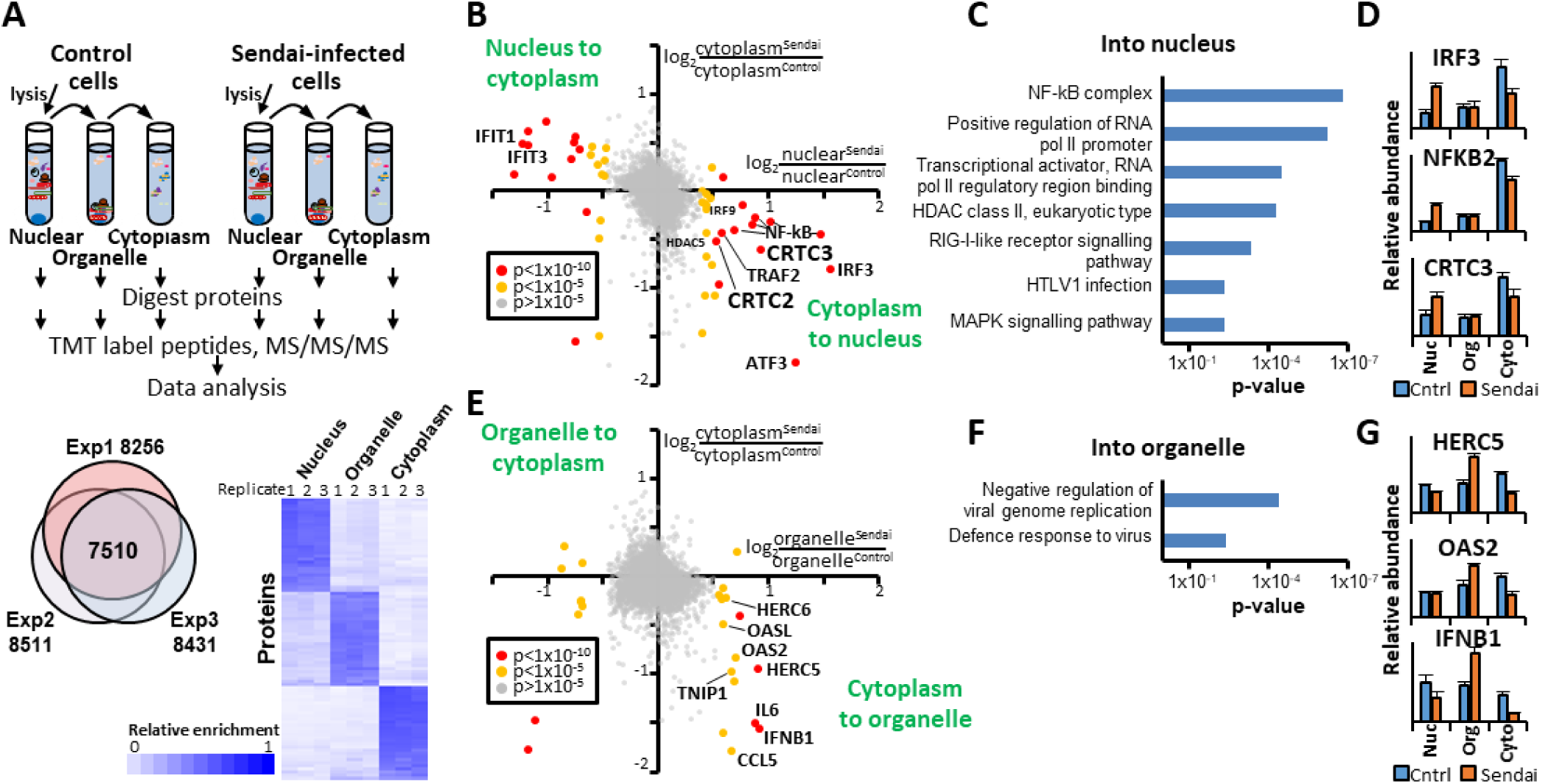
Subcellular profiling during Sendai infection identifies translocation of CRTC2 and CRTC3 from cytoplasm to nucleus. **(A)** (top) Schematic of the experiment. Differential centrifugation of mechanical cellular lysates separated nuclear and organellar fractions, with the supernatant as the cytoplasmic fraction. (bottom) overlap of proteins quantified in each replicate experiment, and hierarchical clustering analysis of markers of each fraction (**Table S3**). (**B**) Scatterplot of nucleo-cytoplasmic movement of proteins quantified in all three replicates (8h infection with Sendai virus or control, n=3). Benjamini-Hochberg corrected significance A values (*18*) were used to estimate p-values for the nuclear Sendai/nuclear control ratio, and dots are coloured according these values. The fold change for cytoplasmic movement was in some cases small, which may reflect a relatively large initial cytoplasmic protein pool. (**C**) Functional enrichment analysis of proteins moving in to the nucleus with p<1×10^-5^ (see also **Table S2C**). (**D**) Example results. Error bars – SEM. (**E**) Scatterplot of organelle-cytoplasmic movement as described in (B). (**F**) Functional enrichment analysis of proteins moving in to organelles with p<1×10^-5^ (see also **Table S2E**). (**G**) Example results. Error bars – SEM.

**Figure S1.**
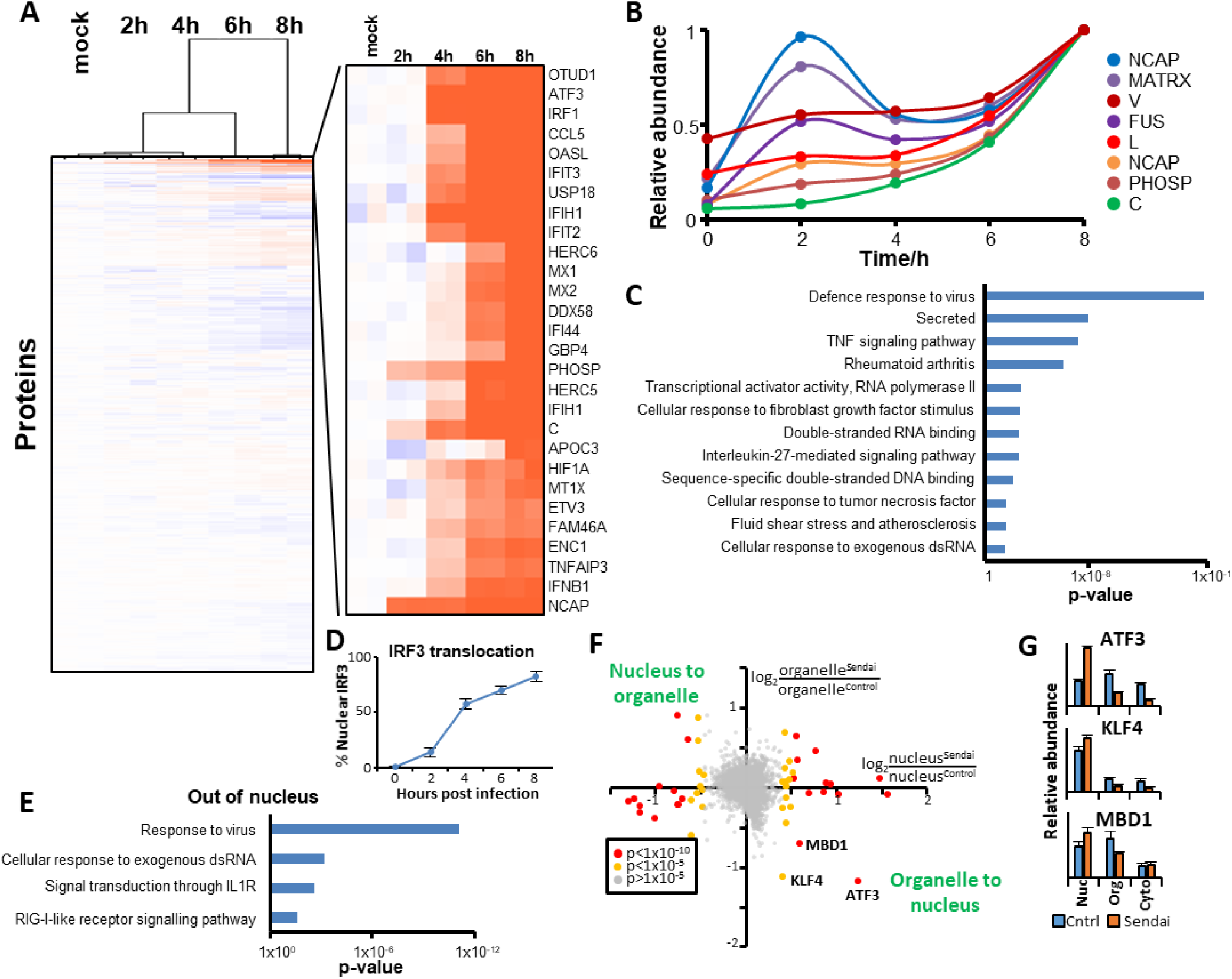
Subcellular and temporal profiling during Sendai infection. (**A**) Hierarchical cluster analysis of 8548 proteins quantified in a temporal proteomic analysis of infection of HFFF-TERTs with Sendai virus (n=2). An enlargement is shown indicating a selection of proteins that were upregulated during the course of the experiment (see also **Table S1**). The vast majority of detected proteins did not show significant abundance changes at this time point. (**B**) Relative abundance over time of all eight viral proteins. (**C**) Functional enrichment analysis of proteins upregulated >2-fold at any time point during infection (**Table S2A**). (**D**) Translocation of IRF3 into the nucleus upon Sendai infection. (**E**) Functional enrichment analysis of proteins moving out of the nucleus with p<1×10^-5^ (See also Figure 1B**, Table S2D**). (**F**) Scatterplot of nucleo-organellar movement of proteins quantified in all three replicates (8h infection with Sendai virus or control, n=3). Benjamini-Hochberg corrected significance A values (*18*) were used to estimate p-values for the nuclearSendai/nuclearControl ratio, and dots are coloured according these values. (**G**) Example results. Error bars – SEM.

Subcellular profiling quantified 9173 human proteins and 8 Sendai proteins in total, with 7510 human proteins quantified in all three biological replicates (**Figure 1A**). An interactive spreadsheet of all proteomic data in this manuscript is provided in **Table S1**, enabling graphical visualisation of each quantified protein. The locations of known marker proteins from each organelle were discriminated well (**Figure 1A, Table S3**). Redistribution of IRF3 and five Rel-like domain-containing proteins of the NF-**κ**B family validated our approach, and DAVID functional enrichment analysis of proteins moving into the nucleus indicated that a variety of transcriptional regulators and components of known antiviral signalling pathways exhibited redistribution upon infection. CREB Regulated Transcription Coactivators-2 and 3 (CRTC2/3) were both amongst proteins exhibiting the greatest degree of cytoplasmic-nuclear translocation, and have not previously been identified in the response to viral infection (**Figures 1B-D, Table S2C**). Proteins redistributing from cytoplasm to nucleus also included the class IIa histone deacetylases (HDAC) 4, 5 and 7; we have previously shown that HDAC4 and 5 can restrict infection by DNA viruses (*19, 20*). A number of proteins appeared to move out of the nucleus into the cytoplasm (**Figure 1B**, top left). However, 19/22 of these were interferon-stimulated genes (interferome 2.01,(*21*)), and 15/22 were upregulated >1.5 fold during the Sendai virus timecourse analysis (**Table S1**) suggesting that this effect may reflect cytoplasmic protein synthesis instead of movement. Proteins seemingly redistributing from cytoplasm to organelle were also enriched in antiviral functions, including oligoadenylate synthetase 2 (OAS2) and 2’-5’-Oligoadenylate Synthetase Like (OASL), and HECT and RLD domain containing E3 ubiquitin protein ligases 5 and 6 (HERC5/6) (**Figures 1E-G, Table S2E**), although this could again represent *de novo* synthesis.

### Early nucleocytoplasmic translocation of CRTC2 and CRTC3 during infection with diverse viruses and in diverse cell types

We confirmed that both ectopically expressed and endogenous CRTC2 and CRTC3 relocated from cytoplasm to nucleus upon Sendai infection by immunofluorescence microscopy (**Figures 2A-B, S2A**). Cells with translocating CRTC2 largely also translocated CRTC3 (**Figure 2C**). To determine whether our findings were limited to infection of fibroblasts by Sendai virus, or represented a more general response to viral infection, we firstly examined other RNA viruses or equivalent stimuli. CRTC2 and CRTC3 both exhibited nucleocytoplasmic relocation upon lipofection with poly (PIC) (**Figures 2D-E**) and infection with respiratory syncytial virus (RSV) (**Figures 2F, S2A**). Importantly, infection with DNA viruses or equivalent stimuli also led to CRTC2/3 translocation, including calf thymus DNA (CT-DNA), modified vaccinia Ankara (MVA), herpes simplex virus-1 (HSV1) and human cytomegalovirus infection (HCMV) (**Figures 2G, S2B-D**).

**Figure 2.**
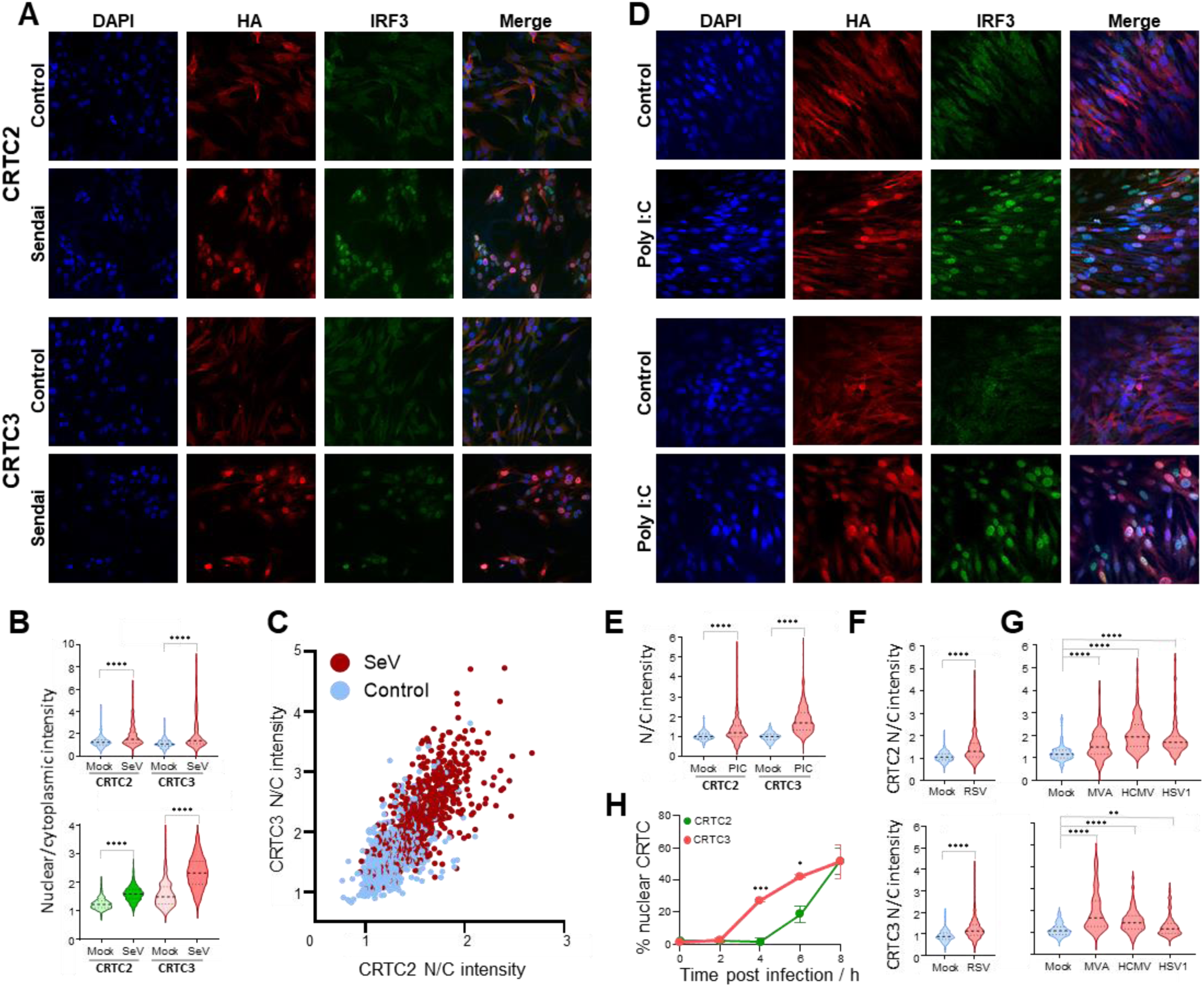
CRTC2 and CRTC3 translocate from cytoplasm to nucleus in response to infection with diverse viruses. Representative immunofluorescence analysis of nucleocytoplasmic movement of HA-tagged CRTC2 and CRTC3, and endogenous IRF3 after 8h of infection with Sendai virus or control. Translocation of endogenous CRTC2/3 is shown in **Figure S3A**. (**B**) Top panel : quantitation of movement shown in (A); bottom panel – quantitation of endogenous CRTC2/3 nuclear/cytoplasmic intensity after 8h of Sendai virus infection or control treatment (representative images shown in **Figure S3A).** CellProfiler was used for quantitation (Methods). Briefly, nuclei were defined by the area of DAPI staining. Cytosol was defined by extending the area of the nuclear mask by 10 pixels then subtracting the area covered by the nuclear mask. A nuclear:cytoplasmic (N:C) ratio was calculated. For this and subsequent subfigures, a minimum of 80 cells were quantified. ****p<0.0001. (**C**) Correlation of CRTC2/3 translocation from cytoplasm to nucleus after Sendai virus infection. Pearson r = 0.7599 (95% confidence interval: 0.7351-0.7826). Two-tailed p-value <0.0001. (**D**) Representative immunofluorescence analysis as (A) 8h post lipofection with poly I:C (4 µg/ml). (**E**) Quantitation of movement shown in (D). ****p<0.0001. (**F**) Quantitation of movement upon RSV infection, from **Figure S2A**. (**G**) Quantitation of movement upon MVA, HCMV and HSV-1 infection, from **Figures S2C-D**. **p<0.01, ****p<0.0001. (**H**) Temporal kinetics of nuclear translocation of stably overexpressed CRTC2 and CRTC3. n=3, counting at least 100 nuclei per replicate over at least two fields. Error bars – SEM. *p<0.05, ***p<0.0005. Scale bar = 100μm.

**Figure S2.**
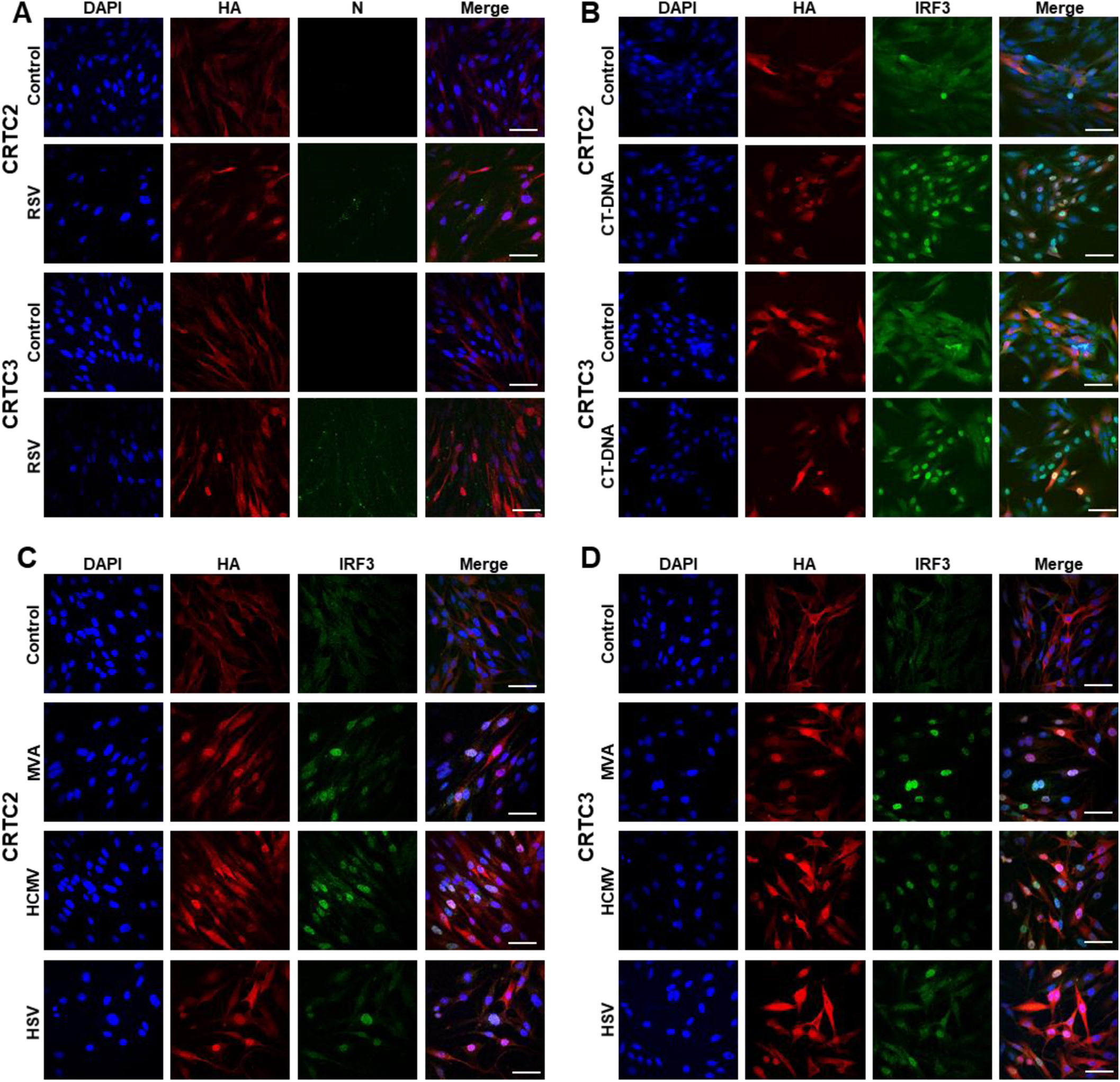
CRTC2 and CRTC3 translocate from cytoplasm to nucleus in response to infection with diverse viruses. Representative immunofluorescence analysis of nucleo-cytoplasmic movement in cells stably expressing HA-tagged CRTC2 or HA-tagged CRTC3. (**A**) after 18h of infection with Respiratory Syncytial virus (RSV) or control (MOI = 2). Green - RSV-N protein. (**B**) after transfection with lipid-delivered Calf Thymus (CT) DNA (0.5μg). (**C-D**) after 6h of infection with Modified Vaccinia Ankara (MVA), Human Cytomegalovirus (HCMV), Herpes Simplex virus 1 (HSV) or control (MOI = 5). Scale bar = 100μm.

CRTC2 also translocated into the nucleus during Sendai virus infection of A549, HUVEC and HBEC3-KT lung epithelial cells (*22*). However, in these cells, CRTC3 exhibited a predominantly nuclear localization prior to infection, although there was a small yet significant increase in nuclear CRTC3 upon infection of HUVEC and A549s (**Figure S3**). To determine the kinetics of CRTC2/3 translocation, we infected HFFF-TERT cells stably expressing HA-tagged CRTC2/3 with Sendai virus. CRTC3 accumulated in nuclei between 2-4 h post infection, whereas CRTC2 did not start to translocate until 4-6 h (**Figure 2H**). The response of CRTC2/3 to multiple stimuli, in diverse cell types suggests that this is an important new pathway in the context of viral infection.

**Figure S3.**
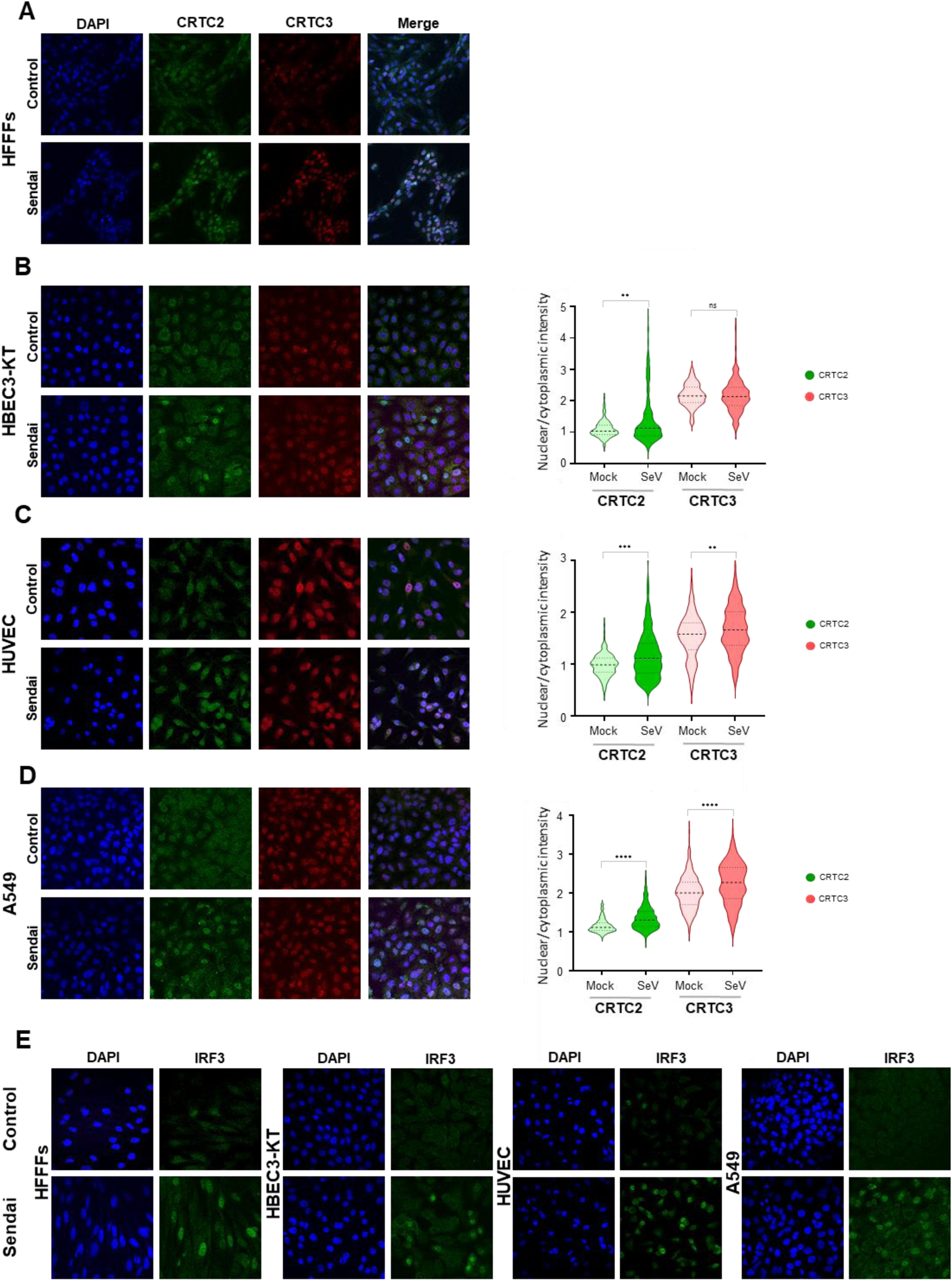
Endogenous CRTC2 and CRTC3 translocate from cytoplasm to nucleus in response to Sendai virus infection in multiple cell types. Representative immunofluorescence analysis of nucleocytoplasmic movement of endogenous CRTC2 and CRTC3 after 8h of infection with Sendai virus (**A**) in HFFF-TERT cells - quantitation is shown in Figure 1C; (**B**) in HBEC3-KT cells; (**C**) in HUVEC/TERT-2 cells; (**D**) in A549 cells. ****p<0.0001. (**E**) Immunofluorescence detection of endogenous IRF3 upon 8h after Sendai infection in HFFF-TERT cells, HBEC3-KT, HUVEC/TERT-2 and A549 cells. Scale bar = 100μm.

### CRTC2 and CRTC3 are required for transcription of a distinct subset of genes in response to viral infection

CRTC2/3 proteins are members of a family of transcriptional co-activators of CREB (cAMP Responsive Element Binding Protein), originally identified as interleukin-8 (IL-8) activators (*23*). CRTC nuclear translocation is an essential, conserved step in activation of a subset cAMP-responsive genes (**Figure 3A**), with key regulatory roles in energy homeostasis, long-term memory and longevity (*24*). CRTC-associated enhancement of CREB dependent transcription appears to be independent of, but synergistic to CREB S133 phosphorylation (*25*).

**Figure 3.**
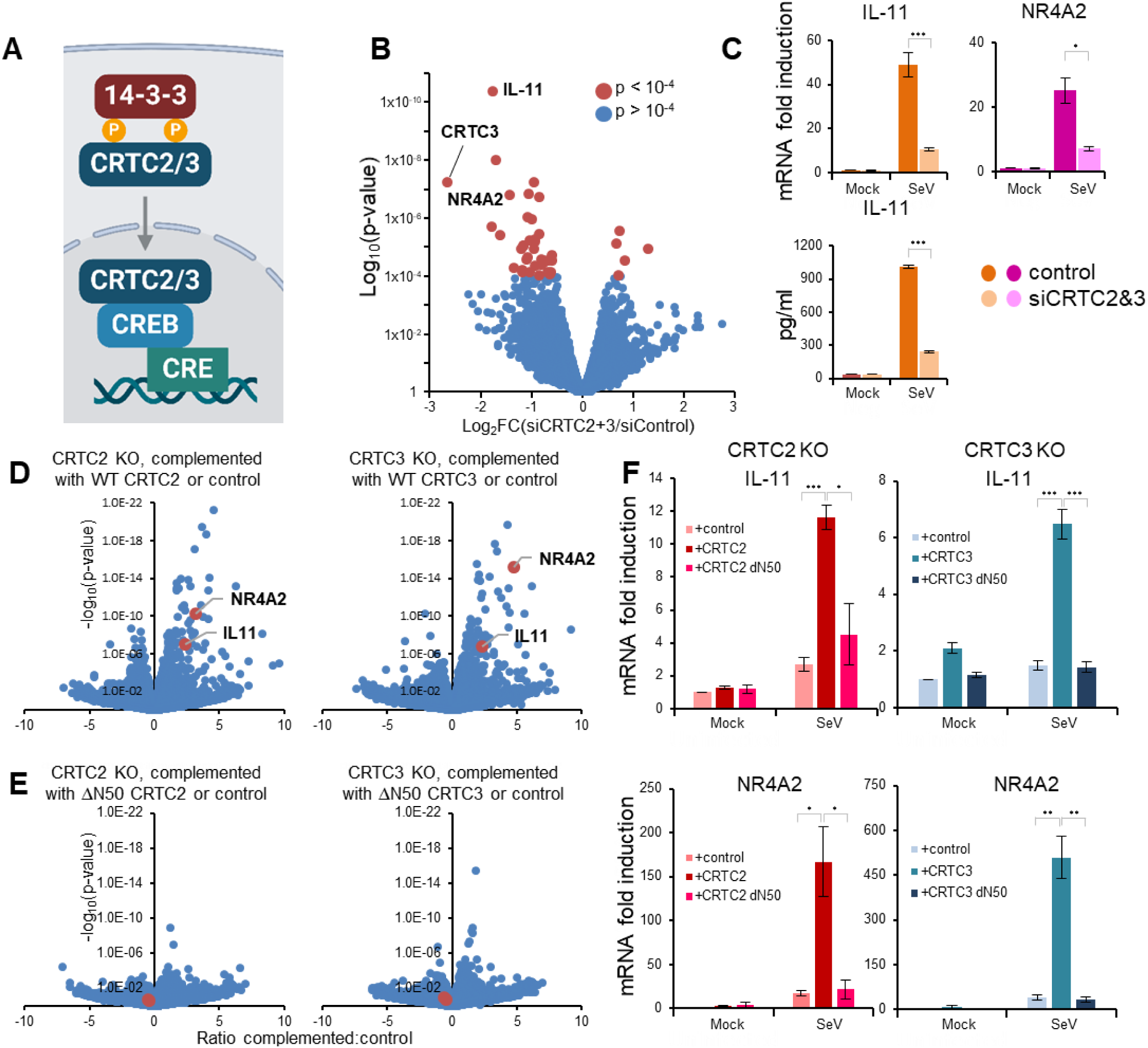
(**A**) Regulation and function of CRTC molecules. Under basal conditions, CRTCs are phosphorylated at two conserved sites and sequestered in the cytoplasm through interactions with 14-3-3 proteins. Dephosphorylated CRTCs translocate to the nucleus, where they bind CREB, stimulating transcriptional activation of cAMP Responsive Element (CRE)-containing target genes. CRTCs all include a CREB-binding domain (CBD), a regulatory region (RD) and a transactivation domain (TAD) (21346730). (**B**) RNAseq comparing HFFF-TERTs knocked down for both CRTC2 and CRTC3 or control then infected with Sendai virus for 8 hours, n=3. CRTC2 was not reliably quantified but depletion was confirmed by qPCR (**Figure S4A**). (**C**) CRTC-dependent induction of IL-11 and NR4A2 in cells knocked down for both CRTC2 and CRTC3 or control then infected with Sendai virus. Top – qPCR analysis for IL-11 (n=4), NR4A2 (n=3). Bottom – ELISA of IL-11 in cellular supernatant. n=4. (**D-E**) Volcano plots showing differential transcript expression by RNAseq in cells singly knocked out for CRTC2 or CRTC3, complemented with the respective wild-type or ΔN50 gene or control then infected with Sendai. n=3. (**F**) qPCR to validate findings in (E) for IL-11 and NR4A2. CRTC2, n=3; CRTC3, n=4. *p<0.05, **p<0.01, ***p<0.001

To determine the effects of CRTC2/3 in the context of viral infection, RNAseq was used to analyse cells depleted of both proteins, then infected with Sendai virus (**Figures 3B, Figure S4A**). The gene most significantly downregulated in the absence of CRTC2/3 was interleukin-11 (IL-11). Interestingly, IL-11 has recently been implicated in fibro-inflammatory processes (*26–29*) and has previously been shown to be released in response to viral infection via an undefined axis (*30–32*). Supporting our results, one of the next most significantly differentially regulated transcripts was Nerve Growth factor IB-like receptor NR4A2 (also known as Nurr1), which has previously been characterized as a CRTC dependent gene and shown to silence HIV infection (*25, 33, 34*). Expression of IL-11 and NR4A2 in a CRTC2- and CRTC3-dependent manner was validated by RT-qPCR and ELISA for IL-11 (**Figure 3C**).

**Figure S4.**
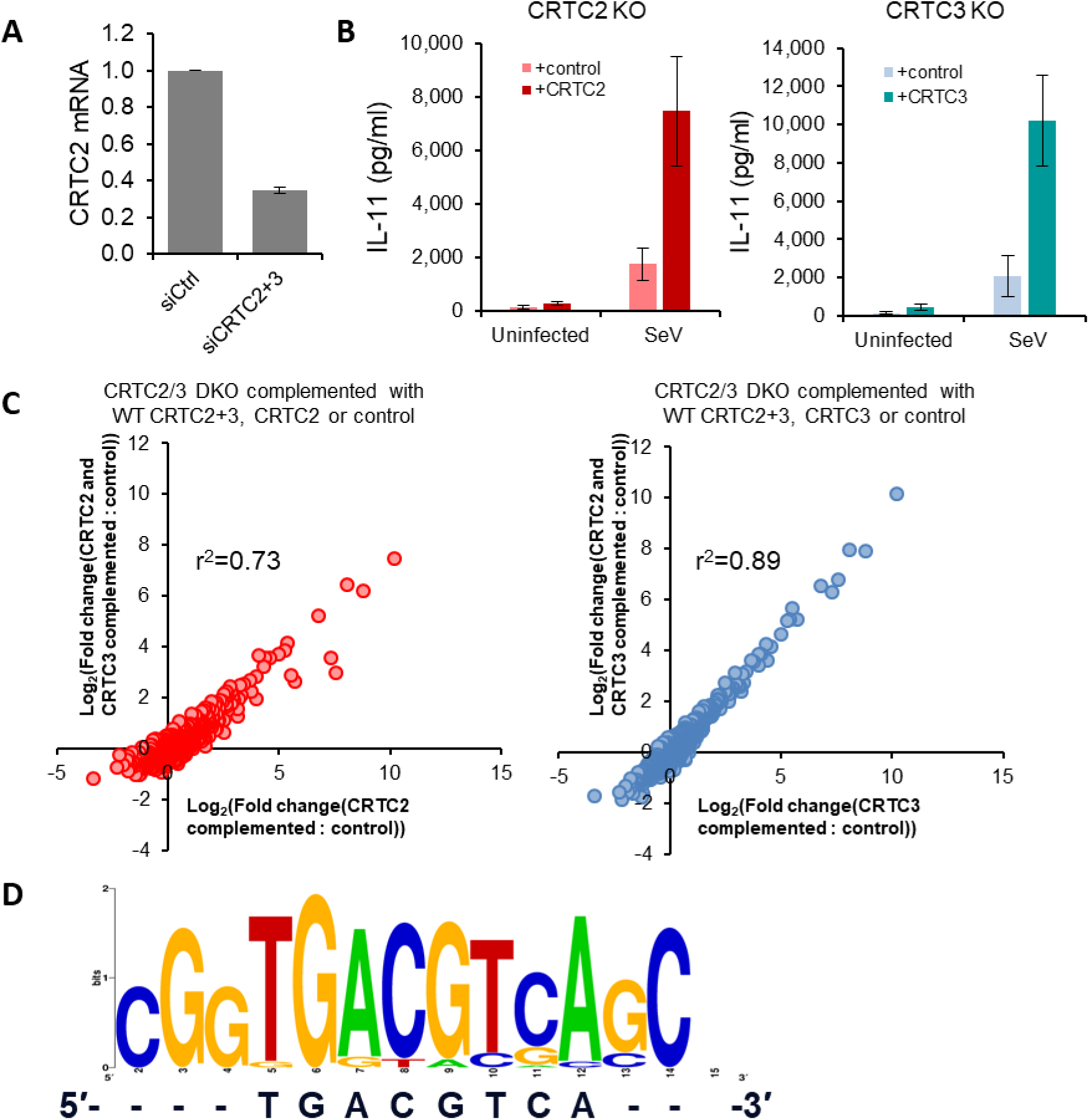
(**A**) qPCR validation of CRTC2 knockdown in cells analysed by RNAseq for Figure 3B. (**B**) ELISA for IL-11 in cellular supernatant of CRTC2 KO (left) or CRTC3 KO cells (right) complemented with the respective wild type protein or mock, then infected with Sendai virus. (**C**) Comparison of transcript abundance in CRTC2/3 DKO cells complemented either with both CRTC2 and CRTC3, or CRTC2 alone (left) or CRTC3 alone (right), or control. In both cases only infected samples are displayed. Transcripts with an average abundance below 10 normalised counts were excluded from downstream analysis. (**D**) Promoter sequences of transcripts enriched in CRTC2/3 DKO HFFF-TERTs complemented either with CRTC2+CRTC3 or control then infected with Sendai virus, compared to the canonical CRE motif (5′-TGACGTCA-3′). The sequences were aligned with Clustal Omega and the sequence logo was generated with WebLogo online tool.

Given that CRTC2 and CRTC3 are from the same family of proteins, we next sought to test whether their role was redundant. Single CRTC2 and CRTC3 knock-out cell lines (CRTC2 KO and CRTC3 KO) in addition to a double knock-out cell line (CRTC2/3 DKO) were generated, then either complemented with the wild type construct, a mutant lacking the N-terminal 50 amino acids (the CREB binding domain for both CRTC2 and CRTC3), or a vector-only control. After infection with Sendai virus, transcripts were sequenced (**Figures 3D-E, S3C, Table S5**).

In CRTC2 KO and CRTC3 KO cells complemented with vector or their respective wild-type gene, IL-11 and NR4A2 were two of the most significantly differentially expressed genes (**Figure 3D**). This occurred despite residual expression of the other CRTC in the single knock-out cells. We suspect that this may be a consequence of overexpression, and note that a replacement of either CRTC2 or CRTC3 alone in CRTC2/3 DKO cells was sufficient to result in a similar transcriptional profile to double-complemented cells (**Figure S4C**). CRTC2/3 function was dependent on the CREB-binding N-terminus of the proteins, as transcription was similar in CRTC2 KO or CRTC3 KO cells complemented with control or the respective ΔN50 CRTC (**Figure 3E**). We confirmed that complementation of either CRTC2 or CRTC3 knockouts with their respective wild-type by not ΔN50 gene was sufficient to enhance transcription of IL-11 and NR4A2 (**Figure 3F**). Together this data strongly suggests that CRTC2 and CRTC3 target the same subset of cellular promoters in response to infection. Analysis of the promoters of CRTC2/3-regulated genes revealed an enrichment of the motif CGGTGACGTCAGC – which contains the CREB1 consensus motif of 5’-TGACGTCA-3’, again supporting the role of CREB in the actions of CRTC2 and CRTC3 (**Figure S4D, Table S4**).

### Nuclear translocation of CRTC2 and CRTC3 during infection is dependent on MAVS, COX2 and protein kinase A

Given the surprising observation that both RNA and DNA stimuli were capable of inducing CRTC2 and CRTC3 nuclear translocation, we attempted to identify the pathway through which this movement was activated. Upon other types of stimuli, CRTC2 and CRTC3 in addition to HDAC4 and 5 (**Figure 1B**) translocate into the nucleus following cAMP/Protein Kinase A (PKA) signalling (*35–37*). We therefore assessed whether the cAMP signalling pathway was activated during Sendai infection. PKA is a canonical effector of cAMP signalling, and immunoblot using a pan-phospho-PKA substrate antibody demonstrated an increase in PKA substrate phosphorylation in Sendai-infected cells, which was inhibited by the PKA inhibitor H89 (**Figures 4A, S5A**).

**Figure 4.**
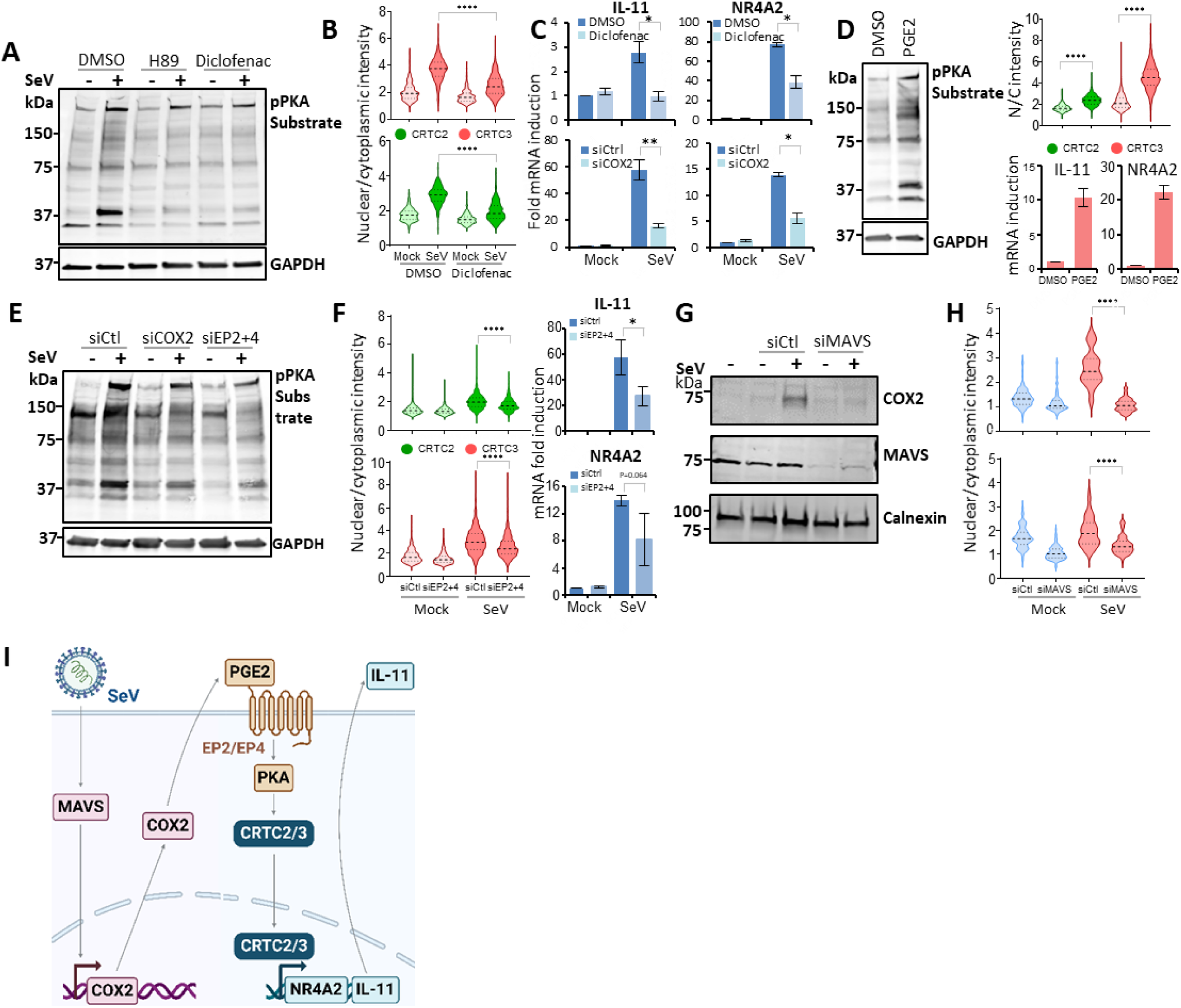
Virus-induced translocation of CRTC2 and CRTC3 is MAVS, COX2- and PKA-dependent. (**A**) Immunoblot showing an increase in PKA substrate phosphorylation following Sendai virus infection. HFFF-TERTs were treated with DMSO, PKA inhibitor H89 (20μM), and COX1/COX2 inhibitor diclofenac (20μM) for the final six hours of an 8 h infection with Sendai virus. Pan-phospho-PKA (pPKA) substrate and GAPDH were analysed; quantitation is shown in Figure S5A. (**B**) Reduced nuclear translocation of endogenous CRTC2 and CRTC3 upon COX inhibition by diclofenac during Sendai virus infection of HFFF-TERTs. For this and subsequent similar subfigures, a minimum of 200 cells were quantified in each of n=3 experiments as described in Figure 2B and Methods. ****p<0.0001. Data is representative of three experiments. (**C**) Reduced transcription of IL-11 and NR4A2 in Sendai virus-infected cells following depletion of COX2 or treatment with diclofenac (20μM). n=3 (IL-11) or 4 (NR4A2). Plotted values represent average +/-SEM. (**D**) Representative immunoblot (n=3), quantitative immunofluorescence (n=3) and qPCR (n=3) demonstrating that PGE2 (20μM) stimulates activation of PKA activation and nuclear translocation of CRTC2 and CRTC3 in HFFF-TERT cells. Quantitation of the immunoblot is shown in Figure S5A. (**E-F**) Representative immunoblot (n=3), quantitative immunofluorescence (n=3) and qPCR (n=3) demonstrating that siRNA depletion of EP2 and EP4 impairs Sendai virus-induced activation of PKA (E), nuclear translocation of CRTC2 and CRTC3 (F-left panels), and IL-11 and NR4A2 transcription. Plotted values represent average +/- SEM.(F-right panels). HFFF-TERT cells were depleted of either COX2 or both EP2 and EP4 then infected with Sendai virus for 8 h. Quantitation of the immunoblot is shown in Figure S5A. (**G-H**) Depletion of MAVS prevents induction of COX2 in response to Sendai virus infection, and prevents nuclear translocation of CRTC2 and CRTC3. Left: quantification of nuclear:cytoplasmic intensity of stably expressed CRTC2 or CRTC3 72 h after depletion of MAVS or control using siRNA, followed by 8h Sendai infection. Representative immunofluorence images are shown in Figure S5A. (**I**) A proposed model for CRTC2/3 translocation in the context of viral infection.

**Figure S5.**
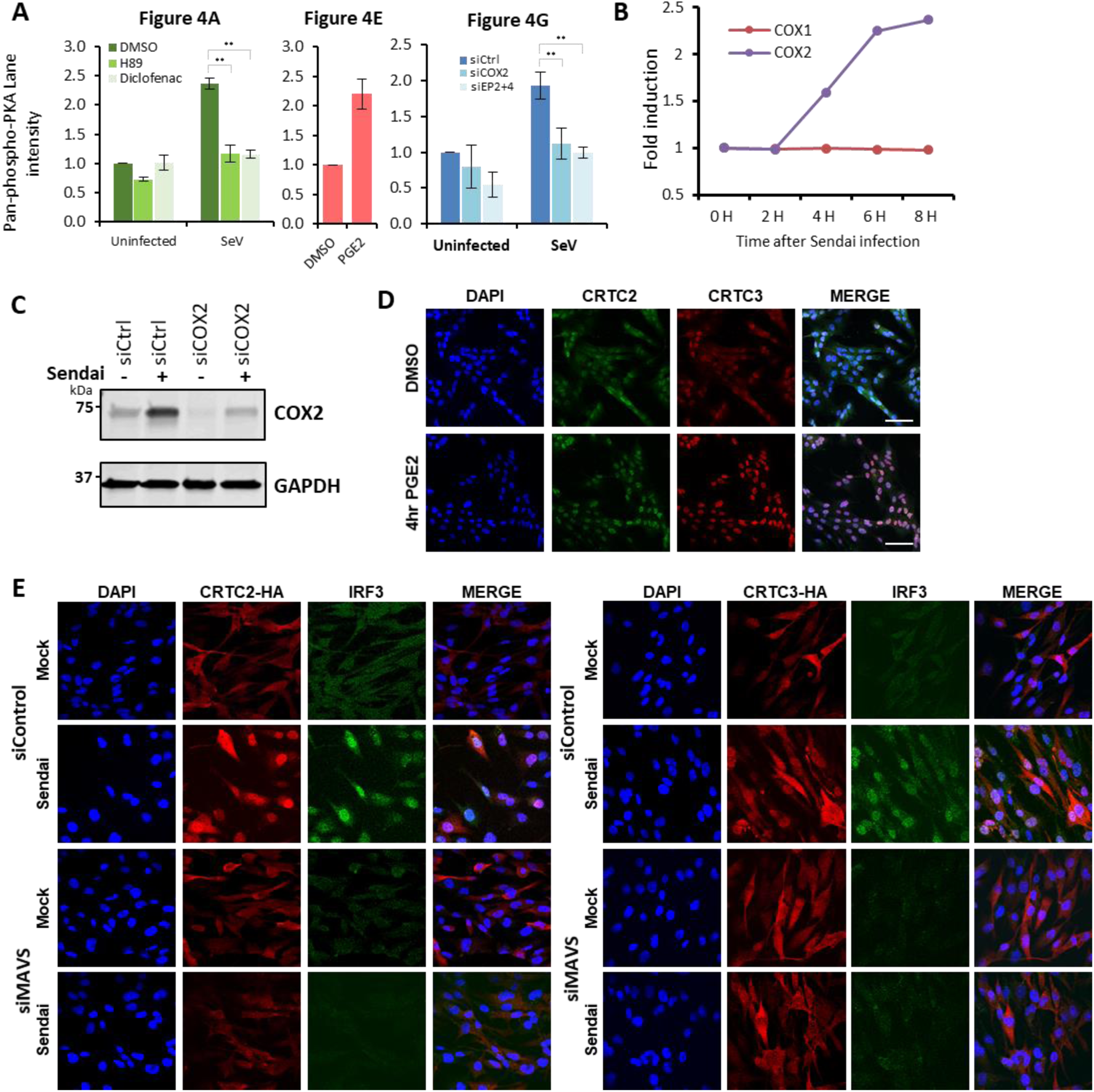
Virus-induced translocation of CRTC2 and CRTC3 is MAVS, COX2- and PKA-dependent. (**A**) Quantitation of pan-phospho-PKA for immunoblots shows in Figures 4A, E and G (n=3). Quantitation was adjusted according to GAPDH intensity for each replicate. **p<0.01, two-tailed Student’s t-test. Error bars – SEM. (**B**) Induction of COX1 and COX2 in HFFF-TERTs infected with Sendai virus, compared to uninfected cells from proteomic analysis. Full data is shown in Table S1. (**C**) Immunoblot demonstrating si-knockdown of COX2 or control in wild type HFFF-TERTs before or after Sendai virus infection for 8h. (**D**) Representative immunofluorescence analysis of nuclear/cytoplasmic movement of endogenous CRTC2 or CRTC3 in wild type HFFF-TERTs treated with DMSO or PGE2 for 4h. (**E**) Representative immunofluorescence images of cells stably expressing HA-tagged CRTC2 or CRTC3 transfected with siRNA control and siRNA targeting MAVS for 72h, followed by 8h Sendai infection. Quantitation of nuclear:cytoplasmic intensity is shown in Figure 4H. Scale bar = 100μm

Many cell surface receptors are coupled to adenylyl cyclase, but relatively few are linked to infection or inflammation. Conversely, elevation of cyclo-oxygenase 2 (COX2) is a well-established component of the inflammatory response which is induced in infection (*38, 39*). This results in elevated levels of prostaglandins driving pyrexia and inflammation (*40, 41*). Our proteomic analysis demonstrated that COX2 but not COX1 was induced in response to Sendai (**Figure S5B**). Blocking both COX1 and COX2 with the non-steroidal anti-inflammatory drug diclofenac resulted not only in a reduction in PKA activity in the context of Sendai virus infection, but also significantly reduced nuclear translocation of CRTC2 and CRTC3, and significantly reduced transcription of IL-11 and NR4A2 (**Figures 4A-C**). Knockdown of COX2 also reduced transcription of IL-11 and NR4A2 (**Figures 4C, S5C**).

Cyclo-oxygenases catalyse production of prostaglandins, and Prostaglandin E2 (PGE2) has previously been shown to induce translocation of CRTC proteins (*42*). We confirmed that PGE2 treatment induced PKA activation, CRTC2 and CRTC3 translocation and transcription of IL-11 and NR4A2 in our system (**Figures 4D, S5D)**. PGE2 signals via four known receptors, EP1-4, of which EP2 and EP4 signal via the adenylyl-cyclase-PKA pathway (*43*) and are expressed in HFFF-TERTs from our RNAseq data (**Table S5**). Depletion of EP2 and EP4 by siRNA was sufficient to impair Sendai virus infection induced activation of PKA, nuclear translocation of CRTC2 and CRTC3 and IL-11 and NR4A2 transcription (**Figures 4E-F**).

MAVS is a core component of the cellular response to RNA virus infection, and is required for downstream activation of IRF3 and NF-κB (*4*). As the COX2 promoter contains an NF-κB response element (*44*), we tested whether MAVS depletion could similarly impair this pathway. Knockdown of MAVS led to a failure to induce COX2 at the protein level in response to Sendai virus infection, and similarly led to a defect in CRTC2/3 nuclear import (**Figures 4G-H, S5E**). A model of this pathway based on the evidence presented in this section is shown in **Figure 4I**.

### Transcriptional upregulation of IL-11 in PBMC from patients with severe COVID-19

Several viral infections are known to induce pulmonary fibrosis, including SARS-CoV-2, Influenza and human Cytomegalovirus (*45–47*). Pulmonary fibrosis frequently complicates COVID-19, although the pathogenesis is unclear. 2-6% of patients with moderate COVID-19 develop fibrosis (*48*), rising to 20-72% if supplemental oxygen or ventilatory support is required (*49*). Since these changes can be irreversible, new predictive tests and treatments are urgently required.

IL-11 is increased in many fibrotic diseases (*50–52*), and blocking IL-11 with neutralising monoclonal antibodies can prevent end organ damage (*53*). However, to our knowledge, there is little available information on a possible role for IL-11 driving COVID-19-associated pulmonary fibrosis, which might be secreted secondary to nuclear translocation of CRTC2/3 after viral sensing. One important study compared peripheral blood mononuclear cell (PBMC) transcriptional profiles from individuals with severe COVID-19 requiring intensive care to intensive care patients without SARS-CoV-2 (*54*) and people with mild COVID-19. We were therefore able to test the hypothesis that IL-11 is raised in severe COVID-19. Interestingly, IL-11 transcripts were significantly higher in PBMC from patients with severe COVID-19 compared to non-SARS-CoV-2 intensive care controls (**Figure 5**). IL-11 transcripts were similarly higher albeit without significance in PBMC from patients with mild COVID-19 compared to uninfected controls.

**Figure 5.**
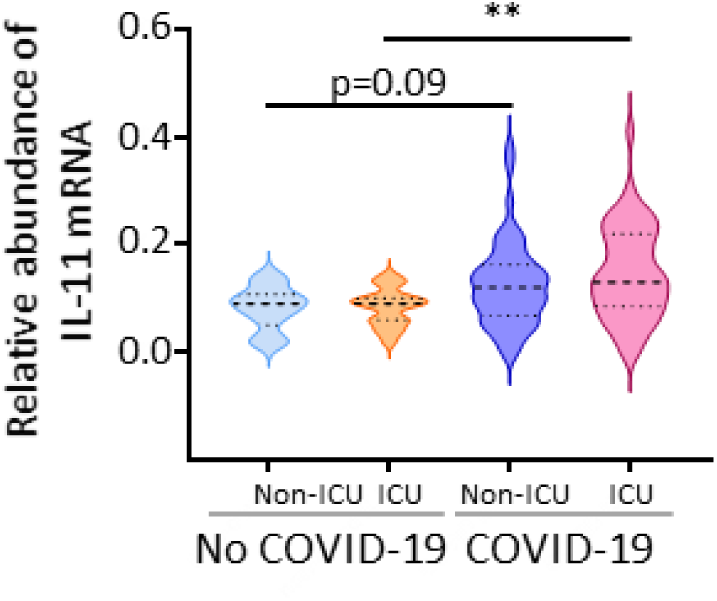
Upregulation of IL-11 transcript in PBMC from individuals with COVID-19 on intensive care, but not intensive care controls. Quantitation of IL-11 mRNA in 10 individuals without COVID-19 not admitted to ICU, 16 non-COVID-19 admitted to ICU, 50 with COVID-19 not admitted to ICU and 50 individuals with COVID-19 admitted to ICU. Data extracted from Shaath & Alajez 2021 (*54*). p-values were estimated using a Mann-Whitney test. **p<0.005.

## DISCUSSION

The detection of pathogens and danger signals through cellular sensors is a defining principle of innate immunity. Although certain core components of the cellular response to viral infection have been extensively studied, systematic unbiased studies of protein translocation offer the potential to identify novel signalling molecules (*14, 55, 56*). Here we identify CRTC2 and CRTC3 as components of the response to viral infection in a pathway downstream of MAVS that is reliant on the production of PGE2, highlighting IL-11 as a secreted cytokine dependent on these coactivators.

The CREB-regulated transcriptional coactivators (also known as the TORC proteins) are a family of structurally related CREB binding proteins which are held inactive in the cytosol by a phosphorylation-dependent interaction with 14-3-3 family proteins (**Figure 3A)**. Upon stimulation, they are dephosphorylated and translocate to the nucleus where they enhance the transcriptional action of CREB in a manner independent of the phosphorylation state of CREB (*25, 35, 57*). Our data demonstrate that transcriptional targets are shared between CRTC2 and CRTC3. This is consistent with previous structural studies of CRTC2 that showed that although CRTC2 makes some contact with DNA, this is for recognition of the minor grove rather than sequence specific base binding (*58*). The fact that CRTC2 has this groove specificity may explain the differential dependence of promoters on CREB and the CRTCs. Similar structural studies of CRTC3 in complex with DNA have not yet been performed. The human genome encodes CRTC1, 2 and 3, although expression varies by tissue type. For example, CRTC1 was quantified very poorly in HFFF-TERT cells, whereas cells in the central nervous system are reported to express high levels of CRTC1 (*35, 59*). Although all three bind to CREB via their N-terminal CREB binding domains, differentiation of function may occur at the level of activation. CRTC1 and CRTC2 have previously been characterised as being released from their 14-3-3 cytosolic partners following the action of the phosphatase calcineurin, whereas CRTC3 relies on dephosphorylation by PP2A following a distinct phosphorylation event, by MAP-kinases or CDKs (*60*). This differential activation may explain the difference in translocation kinetics seen in this study, and potentially provide an evolutionary rationale for coactivators targeting the same subset of genes, but in response to distinct triggers. Given our observation of a role for CRTC2 and 3 in an inflammatory context it is interesting to note that CRTC3 and the adenylyl cyclase-linked PGE2 receptor EP4 have previously been linked to inflammatory bowel disease by genome wide association studies, as has IL-10, which is known to be a CRTC3 transcriptional target in certain cell types (*61, 62*).

IL-11 has hitherto been an under-characterised cytokine, with differing reports on its role in haematopoiesis (*63–66*). IL-11 signals via the IL-11 receptor in complex with the gp130 cell surface receptor which it shares with receptors for IL-6, IL-27, OSM, LIF, CNF, CT-1 and CLCF1. Signalling via IL-11 has recently been implicated in release of inflammatory cytokines such as TNF-α, CCL2 and CCL5 in the context of diet induced steatohepatitis, cardiac fibrosis and experimentally induced lung fibrosis (*50–52*). Although IL-11 release has been previously observed to be released in the context of viral infection (*30–32*), or induced by IL-1B (*67*), its role at the organism level in the context of viral infection requires further research.

Although this work sheds light on a previously unappreciated signalling pathway in viral infection, we acknowledge the limitations of this study. Primarily, our work was *in vitro*, and although our findings fit with an evolving model from the published literature, *in vivo* studies will be required to confirm a role for the CRTCs and IL-11 in viral pathology. It is fascinating to note that IL-11 transcripts were higher in peripheral blood mononuclear cells from patients in intensive care units than non-SARS-CoV-2 infected intensive care controls, particularly as SARS-CoV-2 infection has been associated with lung fibrosis in over 70% of those requiring mechanical ventilation (*49*). This may indicate a role for CRTC-mediated viral signalling in the pathogenesis of this important disease. Further research in this area is clearly warranted, particularly given the existence of IL-11 neutralising monoclonal antibodies, which have been shown to reverse IL-11 induced organ damage and fibrosis in other experimental pathological settings (*50, 68, 69*).

Here we identify CRTC2 and CRTC3 as previously unacknowledged components of the response to viral infection, characterise components required for their nuclear import, and identify a subset of genes upregulated in viral infection in a CRTC-dependent manner. We observe a role for prostaglandins in the movement of both CRTC2 and CRTC3 into the nucleus, and as such it will be interesting to assess the impact of commonly used non-steroidal anti-inflammatories such as ibuprofen on circulating IL-11. This study highlights how spatial proteomics enables systematic mapping of immunologically relevant, new viral sensing pathways.

## ACKNOWLEDGEMENTS

We are grateful to Prof. Steve Gygi for providing access to the “MassPike” software pipeline for quantitative proteomics. We are also grateful to the Cambridge Institute for Medical Research core facilities in flow cytometry and microscopy. This work was supported by a Medical Research Council Project Grant (MR/W025647/1) to MPW, an Addenbrooke’s Charitable Trust Grant (900408) to MPW, an Evelyn Trust Fellowship (project reference 18/27) to BJR, a Medical Research Council Fellowship and BBSRC project grants to BYWC (MR/R021821/1, BB/X001261/1, BB/V017780/1 and BB/V006096/1), and the NIHR Cambridge Biomedical Research Centre (NIHR203312). GHHB was supported by the Max-Planck Society for the Advancement of Science. The views expressed are those of the authors and not necessarily those of the NIHR or the Department of Health and Social care. For the purpose of open access, the authors have applied a CC-BY public copyright license to any Author Accepted Manuscript version arising from this submission.

## AUTHOR CONTRIBUTIONS

Conceptualization: M.P.W. Investigation: B.J.R., M.O., G.W., Y.D., C.D., Y.L., R.A., G.E., N.I., D.J.H., P.L, B.C., G.B. Data analysis: B.J.R., M.O., G.W., M.P.W. Funding acquisition: B.J.R., B.C., M.P.W. Supervision: B.C., G.B., M.P.W. Writing: B.J.R., M.O., G.W., M.P.W.

All authors edited the manuscript.

## MATERIALS AND METHODS

### Cells and cell culture

Human foetal foreskin fibroblast cells immortalised with human telomerase (HFFF-TERTs, male), HEK293T cells (female) and A549 cells were grown in Dulbecco’s modified Eagle’s medium (DMEM) supplemented with foetal bovine serum (FBS: 10% v/v), and 100 IU/ml penicillin / 0.1 mg/ml streptomycin (DMEM/FBS/PS) at 37°C in 5% v/v CO_2_. HFFF-TERTs have been tested at regular intervals since isolation to confirm that human leukocyte antigen (HLA) and MHC Class I Polypeptide-Related Sequence A (MICA) genotypes, cell morphology and antibiotic resistance are unchanged. HEK293T cells were obtained as a gift from Professor Paul Lehner (Cambridge Institute for Therapeutic Immunology and Infectious Disease, Cambridge, UK) and had been authenticated by Short Tandem Repeat profiling (*70*). A549 cells were was isolated from the lung tissue of a white male with lung cancer. HBEC3-KT cells are a hTERT-immortalized lung epithelial cell line, isolated from the bronchus of a female patient. The cell line was acquired from ATCC (CRL-4051). HBEC3-KT cells were grown in Keratinocyte SFM (1X) complemented with 2.5 µg human recombinant epidermal growth factor (rEGF) and 25 mg bovine pituitary extract (BPE). HUVEC/TERT-2 cells (CRL-4053) are an hTERT-immortalized endothelial cell isolated from the vascular endothelium of a female patient. The cells were grown in Human Endothelial Serum Free Medium (SFM) supplemented with 50xGibco™ Large Vessel Endothelial Supplement (LVES). HUVEC/TERT 2 were a gift from Dr. Jeanne Salje (Cambridge Institute for Medical Research, Cambridge, UK), and were acquired from ATCC. All cells were confirmed to be mycoplasma-negative (Lonza MycoAlert).

### Viruses and cellular stimuli

Sendai virus (Cantell strain) was obtained from ATCC (VR-907). This strain is produced in embryonated eggs and produces a high proportion of ‘defective interfering’ genome intermediates in mammalian cells, and little to no productive virus. As such, viral effective titre was determined by the ability to cause IRF3 nuclear translocation for each batch, as ‘MOI’ is difficult to ascertain without a full cycle of infection. A standard dilution of 1:100 Sendai virus to culture medium (2.5% FCS) was used for infections. After 2 hours of infection, supernatant was removed and replaced with full culture medium (10% FCS).

The genome sequence of HCMV strain Merlin is designated the reference for HCMV by the National Center for Biotechnology Information, and was originally sequenced after 3 passages in human fibroblast cells (*71*). A recombinant version (RCMV1111) of this strain was derived by transfection of a sequenced BAC clone (*72*). RCMV1111 contains point mutations in two genes (RL13 and UL128) that enhance replication in fibroblasts (*72*). Viral stocks were prepared from HFFF-TERTs as described previously (*73*). When complete cytopathic effect was observed, cell culture supernatants were centrifuged to remove cell debris and then centrifuged at 22,000 × g for 2 h to pellet cell-free virus. The virus was resuspended in fresh DMEM, and residual debris was removed by centrifugation at 16,000 x g for 1 min.

Modified vaccinia Ankara (MVA) was obtained as a seed stock prepared from an original MVA stock at passage 575 (*74*). MVA was propagated in CEFs, purified by ultracentrifugation through two 36% (w/v) sucrose cushions and suspended in 10 mM Tris-HCl pH 9.0. MVA infectivity was determined by immunocytochemistry on CEFs and HFFF-TERTs cells, by using a polyclonal rabbit anti-VACV antibody (*75*).

A fluorescent HSV-1 strain encoding UL47 with an in-frame C-terminal fusion to EYFP (A206K) was constructed using bacterial artificial chromosome (BAC)-cloned KOS strain of HSV-1 (*76*). Viral stocks were prepared in Vero cells and titrated using a plaque assay on HFFF-TERT monolayers.

Polyinosinic-polycytidylic acid (poly(I:C), Invivogen) was diluted in nuclease-free water (Ambion, ThermoFisher). Calf thymus DNA (CT-DNA) was purchased from Millipore, Merck, and diluted in nuclease-free water (Ambion, ThermoFisher).

### Plasmid construction

For exogenous gene expression, plasmids were constructed by Gateway cloning into the a pHAGE vector under the control of a SFFV promoter (*17*). The pDONR223 was employed as an entry clone as per the manufacturer’s instructions, and as described in Nightingale et al. (*17*). A blasticidin (BSD) resistant pHAGE vector was constructed by subcloning the BSD gene in place of the parental pHAGE puromycin resistance gene. Construct sequences were validated by Sanger or Nanopore sequencing. Inserts were generated by PCR amplification from HFFF-TERT cDNA libraries or in the case of CRTC2 from a plasmid originating with the Harvard Medical School plasmid repository. The ΔN50 CRTC2 and ΔN50 CRTC3 constructs were created by PCR amplification from their respective wild-type parental plasmids.

### Stable cell lines

Stable HFFF-TERT cell lines were generated by lentiviral transduction. HEK293T cells were transfected using TransIT293 as per the manufacturer’s protocol with lentiviral vector, pCMV8.91 and VSVG expression plasmid. 48hrs post transfection lentivirus containing supernatant was collected, and clarified by high speed centrifugation. Supernatant was then either used immediately or stored at -20 °C for later use. Target HFFF-TERT cells were then transduced by application of lentivirus at a range of dilutions, and after approximately 72 hours selected using 1μM puromycin or 1μM blastomycin as appropriate until control untransduced cells were all dead. Dilutions resulting in approximately 1/3 of cells surviving selection were taken forwards for experiments.

### Immunoblotting

Cell samples were lysed in 1x RIPA buffer with protease inhibitors at 4 °C. Lysates were clarified by centrifugation if required. Samples were mixed with 6x Laemmli buffer, boiled for 5 minutes then run on Biorad precast gels (4-15%) in 1x running buffer for 90 minutes at 100 V. Samples were then transferred to PVDF membrane at a constant 350mA for 1 hour. Membranes were then blocked with 5% Marvel solution in Tris-buffered saline with 0.1% Tween 20 detergent (TBST). Primary antibody incubations were routinely performed overnight at 4 °C. After washing, LICOR secondary antibodies were added at 1:10,000 for 1 h while protected from light. Samples were washed again prior to imaging using a LICOR Odyssey CLx instrument.

### Immunofluorescence

Cells were fixed using BD Cytofix fixation buffer as per the manufacturer’s recommendations. Fixed cells were then washed in PBS and stored at 4 °C until used. Cells were permeabilized using ice-cold methanol and then blocked in a solution of 2% FCS in PBS. Subsequent immunostaining was performed in a buffer of PBS, 2% FCS and 0.1% Tween 20 detergent. Cells were mounted in medium containing DAPI and either analysed immediately or stored at 4 °C until analysed. Microscopy was performed on a Zeiss 710, EVOS cell imaging system or Zeiss 880.

### RNA extraction, cDNA and qPCR

RNA was extracted from 5×10^5^ HFFF-TERT cells using the QIAGEN RNeasy kit as per the manufacturer’s protocol. If the extracted RNA was to be used for RNAseq then as part of this protocol then DNA was digested on-column. If RNA was to be used for qPCR then DNase treatment was performed using Invitrogen TURBO DNase as per the manufacturer’s protocol. cDNA was synthesised using Promega GoScript reverse transcriptae kit as per the manufacturer’s instructions, following DNase treatment. Subsequent qPCR was performed using an applied biosystems Fast SYBR Green master mix system, and analysed on a BioRad CFX touch thermocycler. Transcript expression was normalised to GAPDH, and analysis performed in BioRad CFX Maestro software, or by delta-Ct analysis in Microsoft Excel.

### Sythego Cas9/CRISPR

Guide RNA (gRNA) pools were purchased from Synthego as lyophilised RNA. These were resuspended in Tris-EDTA buffer before dilution in molecular biology grade water. gRNAs were then combined with an appropriate ratio of Cas9 protein and nucleofection supplement to form ribonucleoprotein complexes. 1.5×10^5^ HFFF-TERT cells were nucleofected per reaction, using the Lonza 4D-X core unit. Single cell clones were isolated by limiting dilution and knockout verified by Western blot.

### Drugs & Ligands

PGE2 was obtained from Tocris biotechnology (#2296) and resuspended in sterile DMSO. PGE2 was used in experiments at 20μM. Diclofenac was obtained from MedChemExpress (#HY-15036) and resuspended in sterile DMSO. Diclofenac was used in experiments at 20μM. H89 was obtained from LKT laboratories (H0003) and resuspended in sterile DMSO. H89 was used in experiments at 20μM.

For experiments in which drugs were applied during Sendai virus infection, to minimise the risk of interference with cellular entry of the virus, infection was first allowed to occur for two hours. Media were then changed to include this indicated drugs for the remainder of the infection.

### siRNAs

siRNA pools were obtained from Dharmacon as ON-TARGET plus siRNA SMARTpools (CRTC2: L-018947-00-0005; CRTC3: L-014210-01-0005; MAVS: L-024237-00-0005; COX2 (PTGS2): L-004557-00-0005; EP2 (PTGER2): L-005712-00-0005; EP4 (PTGER4): L-005714-00-0005; Non-targeting Control Pool: D-001810-10-05), resuspended as per the manufacturer’s instructions, and stored in aliquots at -80C. HFFF-TERT cells were seeded the day before siRNA transfection. The next day siRNAs were transfected using the transfection reagent DharmaFECT 1 as per the manufacturer’s instructions. A final concentration of 50nM for each siRNA was used.

### ELISA

Indicated cells were infected with Sendai virus for 24 hours prior to supernatant harvest. Supernatants were clarified of any cellular debris by centrifugation prior to storage in aliquots at -80 °C. Supernatants were thawed only once then discarded. Anti-IL11 ELISA assays were performed using either Abcam (ab100551) or R&D (D1100) kits as per the manufacturer’s instructions. Colorimetric readings were measured using a Tecan spark platform.

### RNAseq

Total RNA was extracted by phenol chloroform and poly(A) selective mRNA 3’ end sequencing libraries was prepared using QuantSeq 3’ mRNA-Seq Library Prep Kit-FWD (Lexogen). Libraries were sequenced by Cambridge Genomic Services. Reads were first trimmed to remove adaptors using BBDuk (*77*), before alignment to concatenated human (GRCh38) and SeV (GenBank: AB855653) genomes using STAR (*78*). Reads aligning to annotated genes were counted using HTSeq (*79*) and genes with fewer than 10 counts were excluded from downstream analysis. Normalisation and differential expression analysis was then performed using edgeR (*80*).

### Subcellular fractionation

HFFF-TERT cells were first either infected with Sendai virus or mock treated. At 8 hours post infection cells were washed in PBS, then incubated with hypotonic lysis buffer (25mM Tris-HCl, 50μM Sucrose, 0.5mM MgCl2, 0.2mM EGTA). Cells were harvested after incubation by scraping, and then lysed using a Dounce homogeniser. Following lysis, sucrose concentration was immediately adjusted to physiological osmolarity by addition of an appropriate volume of hypertonic sucrose buffer (25mM Tris-HCl, 2.5M Sucrose, 0.5mM MgCl2, 0.2mM EGTA). The nuclear fraction was then isolated by centrifugation at 1,000g for 10 minutes, and the post-nuclear lysate was centrifuged at 78,400g for 30 minutes. The resulting pellet represents the organellar fraction, and the supernatant the cytosolic fraction. Nuclear and organellar fractions were resuspended in Guanidine-HEPES buffer (6M Guanidine/50 mM HEPES pH 8.5). The cytosolic fraction was concentrated by Trichloro-acetic acid (TCA) precipitation. Briefly, 100% TCA was added to cytosol sample in a 1:5 ratio and incubated at 4 °C. The sample was then centrifuged at 21,000g for 30 minutes at 4 °C. Protein pellets were then washed in TCA and then acetone before resuspension in Guanidine-HEPES buffer.

Subcellular fractions in Guanidine-HEPES buffer were further processed by reduction by dithiothreitol (DTT) at a final concentration of 5mM, then alkylated using iodoacetamide (15mM final concentration). Excess iodoacetamide was then quenched with an excess of DTT. Samples were then diluted with 200mM HEPES pH 8.5 to a final guanidine concentration of 1.5M. Proteins were then digested with LysC protease at a 1:100 protease : protein ratio. Following LysC digest, samples were further diluted in 200mM HEPES to a final guanidine concentration of 0.5M. Samples were then digested with Trypsin at a 1:100 protease : protein ratio overnight at 37°C. The next day the reaction was quenched by the addition of formic acid to a volume of 5%, centrifuged to remove undigested protein, and peptides processed by C18 solid-phase extraction (SPE, Sep-Pak, Waters) and vacuum-centrifuged to near-dryness.

### Whole cell lysate protein digestion for proteomics

Cells were washed twice with phosphate buffered saline (PBS), and 250 μl Guanidine-HEPES buffer added (6M Guanidine/50 mM HEPES pH 8.5). Cell lifters (Corning) were used to scrape cells in lysis buffer, which was removed to an Eppendorf tube, vortexed extensively then sonicated. Cell debris was removed by centrifuging at 21,000 g for 10 min twice. Dithiothreitol (DTT) was added to half of the sample to a final concentration of 5mM and incubated at room temperature for 20 mins. Cysteine residues were alkylated with 15 mM iodoacetamide and incubated for 20 min at room temperature in the dark. Excess iodoacetamide was quenched with DTT for 15 mins. Samples were diluted with 200 mM HEPES pH 8.5 to 1.5 M Guanidine followed by digestion at room temperature for 3 h with LysC protease at a 1:100 protease-to-protein ratio. Samples were further diluted with 200mM HEPES pH8.5 to 0.5 M Guanidine. Trypsin was then added at a 1:100 protease-to-protein ratio followed by overnight incubation at 37°C. The reaction was quenched with 5% formic acid and centrifuged at 21,000 g for 10 min to remove undigested protein. Peptides were subjected to C18 solid-phase extraction (SPE, Sep-Pak, Waters) and vacuum-centrifuged to near-dryness.

### Peptide labelling with tandem mass tags (TMT)

In preparation for TMT labelling, desalted peptides were dissolved in 200mM HEPES pH8.5. TMT reagents were dissolved in anhydrous acetonitrile and 3 μl added to peptide at a final acetonitrile concentration of 30% (v/v). Following incubation at room temperature for 1 h, the reaction was quenched with hydroxylamine to a final concentration of 0.3% (v/v). TMT-labelled samples were combined at a 1:1:1:1:1:1 (subcellular fractionation) or 1:1:1:1:1:1:1:1:1:1 (whole cell lysate) ratio. The sample was vacuum-centrifuged to near dryness and subjected to C18 SPE (Sep-Pak, Waters). An unfractionated single shot was analysed initially to ensure similar peptide loading across each TMT channel, thus avoiding the need for excessive electronic normalization.

### Offline HpRP fractionation

TMT-labelled tryptic peptides were subjected to HpRP fractionation using an Ultimate 3000 RSLC UHPLC system (Thermo Fisher Scientific) equipped with a 2.1 mm internal diameter (ID) x 25 cm long, 1.7 μm particle Kinetix Evo C18 column (Phenomenex). Mobile phase consisted of A: 3% acetonitrile (MeCN), B: MeCN and C: 200 mM ammonium formate pH 10. Isocratic conditions were 90% A/10% C, and C was maintained at 10% throughout the gradient elution. Separations were conducted at 45 °C. Samples were loaded at 200 μl/min for 5 min. The flow rate was then increased to 400 μl/min over 5 min, after which the gradient elution proceed as follows: 0-19% B over 10 min, 19-34% B over 14.25 min, 34-50% B over 8.75 min, followed by a 10 min wash at 90% B. UV absorbance was monitored at 280 nm and 15 s fractions were collected into 96 well microplates using the integrated fraction collector. Fractions were recombined orthogonally in a checkerboard fashion, combining alternate wells from each column of the plate into a single fraction, and commencing combination of adjacent fractions in alternating rows. Wells were excluded prior to the start or after the cessation of elution of peptide-rich fractions, as identified from the UV trace.

### LC-MS3

MS data were generated using an Orbitrap Fusion Lumos (Thermo). An Ultimate 3000 RSLC UHPLC machine equipped with a 300 μm internal diameter × 5 mm Acclaim PepMap μ-Precolumn (Thermo) and a 75 μm internal dimeter × 50 cm 2.1 μm particle Acclaim PepMap RSLC analytical column were used. The loading solvent was 0.1% FA. The analytical solvent consisted of 0.1% FA (A) and 80% AcN + 0.1% FA (B). All separations were carried out at 40°C. Samples were loaded at 5 μl/min for 5 min in loading solvent. The analytical gradient consisted of 3-7% B over 3 min, 7-37% B over 173 min, followed by a 4 min wash at 95% B and equilibration at 3% B for 15 min. Each analysis used a MultiNotch MS3-based TMT method (*81*). The following settings were used: MS1: 380-1500 Th, 120,000 resolution, 2×10^5^ automatic gain control (AGC) target, 50 ms maximum injection time. MS2: Quadrupole isolation at an isolation width of mass-to-charge ratio (m/z) 0.7, collision-induced dissociation fragmentation (normalized collision energy (NCE) 35) with ion trap scanning in turbo mode from m/z 120, 1.5×10^4^ AGC target, 120 ms maximum injection time. MS3: In Synchronous Precursor Selection mode, the top 10 MS2 ions were selected for higher energy collision dissociation (HCD) fragmentation (NCE 65) and scanned in the Orbitrap at 60,000 resolution with an AGC target of 1×10^5^ and a maximum accumulation time of 150 ms. Ions were not accumulated for all parallelisable time. The entire MS/MS/MS cycle had a target time of 3 s. Dynamic exclusion was set to ± 10 ppm for 70 s. MS2 fragmentation was triggered on precursors 5×10^3^ counts and above.

### Proteomic data analysis

In the following description, the first report in the literature for each relevant algorithm is listed. Mass spectra were processed using MassPike, which is a Sequest-based software pipeline for quantitative proteomics, through a collaborative arrangement with Professor Steven Gygi’s laboratory at Harvard Medical School. MS spectra were converted to mzXML using an extractor built upon Thermo Fisher’s RAW File Reader library (version 4.0.26). This software is a component of the MassPike software platform and is licensed by Harvard Medical School.

A combined database was constructed as described in (*17*)from (a) the human Uniprot database (accessed 26 January 2017), (b) all Sendai virus proteins (c) common contaminants such as porcine trypsin and endoproteinase LysC. The combined database was concatenated with a reverse database composed of all protein sequences in reversed order. Searches were performed using a 20 ppm precursor ion tolerance (*82*) Product ion tolerance was set to 0.03 Th. Oxidation of methionine residues (15.99492 Da) was set as a variable modification. Peptides were assumed to be fully tryptic with up to two missed cleavages.

To control the fraction of erroneous protein identifications, a target-decoy strategy was employed (*83, 84*). Peptide spectral matches (PSMs) were filtered to an initial peptide-level false discovery rate (FDR) of 1% with subsequent filtering to attain a final protein-level FDR of 1% (*85, 86*). PSM filtering was performed using linear discriminant analysis as described previously (*87*). Filtering was implemented in R using the linear discriminant analysis (LDA) function in the package MASS (cran.r-project.org/web/packages/MASS). This distinguishes correct from incorrect peptide identifications in a manner analogous to the widely used Percolator algorithm (*88*) although employing a distinct machine-learning algorithm. The following parameters were considered: XCorr, ΔCn, missed cleavages, peptide length, charge state, and precursor mass accuracy. Peptides shorter than seven amino acids in length or with XCorr less than 1.0 were excluded prior to LDA filtering.

Protein assembly was guided by principles of parsimony to produce the smallest set of proteins necessary to account for all observed peptides (algorithm described in (*87*). Proteins were quantified by summing TMT reporter ion counts across all matching peptide-spectral matches using “MassPike”, as described previously (*81*). Briefly, a 0.003 Th window around the theoretical m/z of each reporter ion was scanned for ions and the maximum intensity nearest to the theoretical m/z was used. The primary determinant of quantitation quality is the number of TMT reporter ions detected in each MS3 spectrum, which is directly proportional to the signal-to-noise (S:N) ratio observed for each ion. An isolation specificity filter with a cut-off of 50% was additionally employed to minimise peptide co-isolation (*81*). Peptide-spectral matches with poor quality MS3 spectra (a combined S:N ratio of less than 250 across all TMT reporter ions) or no MS3 spectra at all were excluded from quantitation. Peptides meeting the stated criteria for reliable quantitation were then summed by parent protein, in effect weighting the contributions of individual peptides to the total protein signal based on their individual TMT reporter ion yields. Protein quantitation values were exported for further analysis in Excel.

For protein quantitation, reverse and contaminant proteins were removed, then each reporter ion channel was summed across all quantified proteins and normalised assuming equal protein loading across all channels. For further analysis and display in Figures, fractional TMT signals were used (i.e. reporting the fraction of maximal signal observed for each protein in each TMT channel, rather than the absolute normalized signal intensity). This effectively corrected for differences in the numbers of peptides observed per protein.

### Pathway analysis

The Database for Annotation, Visualisation and Integrated Discovery (DAVID) was used to determine functional enrichment (*89*). A given cluster was always searched against a background of all proteins quantified.

### Cellprofiler

A nuclear:cytosolic fluorescence ratio was quantified using a custom pipeline in Cellprofiler (*90*). In brief, nuclei were defined on the basis of the area covered by DAPI staining, and a mask created based on this region. Nuclei touching the edge of the field of view were excluded.

To measure the cytoplasmic fluorescence intensity the area of the nuclear mask was extended by 10 pixels, then the area covered by the nuclear mask was subtracted. The remaining area was defined as the cytosol. This approach was chosen as opposed to attempting to quantify the entire cytosol to compensate for differences in cell shape, size and crowding. The nuclear:cytosplasmic ratio was then calculated for each nucleus by dividing the mean fluorescence of each nucleus by the mean fluorescence of the associated cytosolic field. To access the statistical significance of the conditions quantified on CellProfiller, two-tailed p-values using the non-parametric Mann-Whitney test were calculated using Graphpad Prism 9.3.1. The Pearson correlation coefficient (r) and 95% confidence intervals were estimated using Graphpad Prism 9.3.1.

### Sequence alignment

53 promoter sequences were enriched in the RNA-Seq dataset comparing CRTC2/3 DKO HFFF-TERTs complemented either with CRTC2+CRTC3 or control then infected with Sendai virus. The sequences were aligned using the Clustal Omega (https://www.ebi.ac.uk/jdispatcher/msa/clustalo) tool for multiple sequence alignments. The resulting alignment was upload to WebLogo 2.8.2 (https://weblogo.berkeley.edu/) to generate the sequence logo based on the base conservation and frequency.

## Data Availability

The mass spectrometry data raw files will be deposited to the ProteomeXchange Consortium via the PRIDE partner repository (*91*) The RNA sequencing raw data files generated in this study is available via the ArrayExpress database with accession E-MTAB-14001.

## REFERENCES

1. S. M. Horner, H. M. Liu, H. S. Park, J. Briley, M. Gale, Jr., Mitochondrial-associated endoplasmic reticulum membranes (MAM) form innate immune synapses and are targeted by hepatitis C virus. Proc Natl Acad Sci U S A 108, 14590–14595 (2011).

2. T. Saitoh et al., Atg9a controls dsDNA-driven dynamic translocation of STING and the innate immune response. Proc Natl Acad Sci U S A 106, 20842–20846 (2009).

3. K. Mukai et al., Activation of STING requires palmitoylation at the Golgi. Nat Commun 7, 11932 (2016).

4. R. B. Seth, L. Sun, C. K. Ea, Z. J. Chen, Identification and characterization of MAVS, a mitochondrial antiviral signaling protein that activates NF-kappaB and IRF 3. Cell 122, 669–682 (2005).

5. T. Abe, G. N. Barber, Cytosolic-DNA-mediated, STING-dependent proinflammatory gene induction necessitates canonical NF-kappaB activation through TBK1. J Virol 88, 5328–5341 (2014).

6. H. Ishikawa, G. N. Barber, STING is an endoplasmic reticulum adaptor that facilitates innate immune signalling. Nature 455, 674–678 (2008).

7. T. Liu, L. Zhang, D. Joo, S. C. Sun, NF-kappaB signaling in inflammation. Signal Transduct Target Ther 2, 17023- (2017).

8. M. Al Hamrashdi, G. Brady, Regulation of IRF3 activation in human antiviral signaling pathways. Biochem Pharmacol 200, 115026 (2022).

9. J. Wang, P. Li, M. X. Wu, Natural STING Agonist as an “Ideal” Adjuvant for Cutaneous Vaccination. J Invest Dermatol 136, 2183–2191 (2016).

10. J. Wang et al., Pulmonary surfactant-biomimetic nanoparticles potentiate heterosubtypic influenza immunity. Science 367, (2020).

11. M. Motwani, S. Pesiridis, K. A. Fitzgerald, DNA sensing by the cGAS-STING pathway in health and disease. Nat Rev Genet 20, 657–674 (2019).

12. S. Jiao et al., Targeting IRF3 as a YAP agonist therapy against gastric cancer. J Exp Med 215, 699–718 (2018).

13. D. N. Itzhak et al., A mass spectrometry-based approach for mapping protein subcellular localization reveals the spatial proteome of mouse primary neurons. Cell Rep 20, 2706–2718 (2017).

14. D. N. Itzhak, S. Tyanova, J. Cox, G. H. Borner, Global, quantitative and dynamic mapping of protein subcellular localization. eLife 5, (2016).

15. L. Martinez-Gil et al., A Sendai virus-derived RNA agonist of RIG-I as a virus vaccine adjuvant. J Virol 87, 1290–1300 (2013).

16. A. Baum, R. Sachidanandam, A. Garcia-Sastre, Preference of RIG-I for short viral RNA molecules in infected cells revealed by next-generation sequencing. Proc Natl Acad Sci U S A 107, 16303–16308 (2010).

17. K. Nightingale et al., High-definition analysis of host protein stability during Human Cytomegalovirus infection reveals antiviral factors and viral evasion mechanisms. Cell Host Microbe 12, 447–460 (2018).

18. J. Cox, M. Mann, MaxQuant enables high peptide identification rates, individualized ppb-range mass accuracies and proteome-wide protein quantification. Nature Biotechnology 26, 1367–1372 (2008).

19. Y. Lu et al., Histone deacetylase 4 promotes type I interferon signaling, restricts DNA viruses, and is degraded via vaccinia virus protein C6. Proc Natl Acad Sci U S A 116, 11997–12006 (2019).

20. L. Soday et al., Quantitative temporal proteomic analysis of vaccinia virus infection reveals regulation of histone deacetylases by an interferon antagonist. Cell Rep 27, 1920–1933 e1927 (2019).

21. I. Rusinova et al., Interferome v2.0: an updated database of annotated interferon-regulated genes. Nucleic acids research 41, D1040–1046 (2013).

22. R. D. Ramirez et al., Immortalization of human bronchial epithelial cells in the absence of viral oncoproteins. Cancer Res 64, 9027–9034 (2004).

23. V. Iourgenko et al., Identification of a family of cAMP response element-binding protein coactivators by genome-scale functional analysis in mammalian cells. Proc Natl Acad Sci U S A 100, 12147–12152 (2003).

24. J. Tasoulas, L. Rodon, F. J. Kaye, M. Montminy, A. L. Amelio, Adaptive Transcriptional Responses by CRTC Coactivators in Cancer. Trends Cancer 5, 111–127 (2019).

25. M. D. Conkright et al., TORCs: transducers of regulated CREB activity. Mol Cell 12, 413–423 (2003).

26. B. Ng, S. A. Cook, S. Schafer, Interleukin-11 signaling underlies fibrosis, parenchymal dysfunction, and chronic inflammation of the airway. Exp Mol Med 52, 1871–1878 (2020).

27. K. Y. Fung et al., Emerging roles for IL-11 in inflammatory diseases. Cytokine 149, 155750 (2022).

28. M. Seyedsadr et al., IL-11 induces NLRP3 inflammasome activation in monocytes and inflammatory cell migration to the central nervous system. Proc Natl Acad Sci U S A 120, e2221007120 (2023).

29. S. Schafer et al., IL-11 is a crucial determinant of cardiovascular fibrosis. Nature 552, 110–115 (2017).

30. O. Einarsson, G. P. Geba, Z. Zhu, M. Landry, J. A. Elias, Interleukin-11: stimulation in vivo and in vitro by respiratory viruses and induction of airways hyperresponsiveness. J Clin Invest 97, 915–924 (1996).

31. H. Bartz, F. Buning-Pfaue, O. Turkel, U. Schauer, Respiratory syncytial virus induces prostaglandin E2, IL-10 and IL-11 generation in antigen presenting cells. Clin Exp Immunol 129, 438–445 (2002).

32. K. L. R. Gustafsson, T. Renne, C. Soderberg-Naucler, L. M. Butler, Human cytomegalovirus replication induces endothelial cell interleukin-11. Cytokine 111, 563–566 (2018).

33. J. H. Kim et al., CREB coactivators CRTC2 and CRTC3 modulate bone marrow hematopoiesis. Proc Natl Acad Sci U S A 114, 11739–11744 (2017).

34. F. Ye et al., Recruitment of the CoREST transcription repressor complexes by Nerve Growth factor IB-like receptor (Nurr1/NR4A2) mediates silencing of HIV in microglial cells. PLoS Pathog 18, e1010110 (2022).

35. T. Sonntag et al., Analysis of a cAMP regulated coactivator family reveals an alternative phosphorylation motif for AMPK family members. PLoS One 12, e0173013 (2017).

36. E. Kozhemyakina, T. Cohen, T. P. Yao, A. B. Lassar, Parathyroid hormone-related peptide represses chondrocyte hypertrophy through a protein phosphatase 2A/histone deacetylase 4/MEF2 pathway. Mol Cell Biol 29, 5751–5762 (2009).

37. J. L. Belfield, C. Whittaker, M. Z. Cader, S. Chawla, Differential effects of Ca2+ and cAMP on transcription mediated by MEF2D and cAMP-response element-binding protein in hippocampal neurons. The Journal of biological chemistry 281, 27724–27732 (2006).

38. N. S. Kirkby et al., Differential COX-2 induction by viral and bacterial PAMPs: Consequences for cytokine and interferon responses and implications for anti-viral COX-2 directed therapies. Biochem Biophys Res Commun 438, 249–256 (2013).

39. N. N. Rumzhum, A. J. Ammit, Cyclooxygenase 2: its regulation, role and impact in airway inflammation. Clin Exp Allergy 46, 397–410 (2016).

40. A. Blomqvist, D. Engblom, Neural Mechanisms of Inflammation-Induced Fever. Neuroscientist 24, 381–399 (2018).

41. E. Ricciotti, G. A. FitzGerald, Prostaglandins and inflammation. Arteriosclerosis, thrombosis, and vascular biology 31, 986–1000 (2011).

42. K. F. MacKenzie et al., PGE(2) induces macrophage IL-10 production and a regulatory-like phenotype via a protein kinase A-SIK-CRTC3 pathway. Journal of immunology (Baltimore, Md. : 1950) 190, 565–577 (2013).

43. Y. Sugimoto, S. Narumiya, Prostaglandin E receptors. The Journal of biological chemistry 282, 11613–11617 (2007).

44. B. Kaltschmidt, R. A. Linker, J. Deng, C. Kaltschmidt, Cyclooxygenase-2 is a neuronal target gene of NF-kappaB. BMC Mol Biol 3, 16 (2002).

45. V. Ojha, A. Mani, N. N. Pandey, S. Sharma, S. Kumar, CT in coronavirus disease 2019 (COVID-19): a systematic review of chest CT findings in 4410 adult patients. Eur Radiol 30, 6129–6138 (2020).

46. N. Nakajima et al., Histopathological and immunohistochemical findings of 20 autopsy cases with 2009 H1N1 virus infection. Mod Pathol 25, 1–13 (2012).

47. E. Coclite, C. Di Natale, G. Nigro, Congenital and perinatal cytomegalovirus lung infection. J Matern Fetal Neonatal Med 26, 1671–1675 (2013).

48. E. Bazdyrev et al., Lung Fibrosis after COVID-19: Treatment Prospects. Pharmaceuticals (Basel) 14, (2021).

49. C. F. McGroder et al., Pulmonary fibrosis 4 months after COVID-19 is associated with severity of illness and blood leucocyte telomere length. Thorax 76, 1242–1245 (2021).

50. A. A. Widjaja et al., Inhibiting Interleukin 11 Signaling Reduces Hepatocyte Death and Liver Fibrosis, Inflammation, and Steatosis in Mouse Models of Nonalcoholic Steatohepatitis. Gastroenterology 157, 777–792 e714 (2019).

51. M. Sweeney et al., Cardiomyocyte-Restricted Expression of IL11 Causes Cardiac Fibrosis, Inflammation, and Dysfunction. Int J Mol Sci 24, (2023).

52. B. Ng et al., Fibroblast-specific IL11 signaling drives chronic inflammation in murine fibrotic lung disease. FASEB J 34, 11802–11815 (2020).

53. S. A. Cook, S. Schafer, Hiding in Plain Sight: Interleukin-11 Emerges as a Master Regulator of Fibrosis, Tissue Integrity, and Stromal Inflammation. Annu Rev Med 71, 263–276 (2020).

54. H. Shaath, N. M. Alajez, Identification of PBMC-based molecular signature associational with COVID-19 disease severity. Heliyon 7, e06866 (2021).

55. A. K. Davies et al., AP-4-mediated axonal transport controls endocannabinoid production in neurons. Nat Commun 13, 1058 (2022).

56. N. Krahmer et al., Organellar Proteomics and Phospho-Proteomics Reveal Subcellular Reorganization in Diet-Induced Hepatic Steatosis. Dev Cell 47, 205–221 e207 (2018).

57. R. A. Screaton et al., The CREB coactivator TORC2 functions as a calcium- and cAMP-sensitive coincidence detector. Cell 119, 61–74 (2004).

58. Y. Song et al., Structural Insights into the CRTC2-CREB Complex Assembly on CRE. Journal of molecular biology 430, 1926–1939 (2018).

59. J. Y. Altarejos, M. Montminy, CREB and the CRTC co-activators: sensors for hormonal and metabolic signals. Nat Rev Mol Cell Biol 12, 141–151 (2011).

60. T. Sonntag et al., Mitogenic signals stimulate the CREB coactivator CRTC3 through PP2A recruitment. iScience 11, 134–145 (2019).

61. L. Jostins et al., Host-microbe interactions have shaped the genetic architecture of inflammatory bowel disease. Nature 491, 119–124 (2012).

62. M. N. Wein, M. Foretz, D. E. Fisher, R. J. Xavier, H. M. Kronenberg, Salt-Inducible Kinases: Physiology, Regulation by cAMP, and Therapeutic Potential. Trends Endocrinol Metab 29, 723–735 (2018).

63. S. R. Paul et al., Molecular cloning of a cDNA encoding interleukin 11, a stromal cell-derived lymphopoietic and hematopoietic cytokine. Proc Natl Acad Sci U S A 87, 7512–7516 (1990).

64. X. X. Du, T. Neben, S. Goldman, D. A. Williams, Effects of recombinant human interleukin-11 on hematopoietic reconstitution in transplant mice: acceleration of recovery of peripheral blood neutrophils and platelets. Blood 81, 27–34 (1993).

65. H. H. Nandurkar et al., Adult mice with targeted mutation of the interleukin-11 receptor (IL11Ra) display normal hematopoiesis. Blood 90, 2148–2159 (1997).

66. S. A. Cook, Understanding interleukin 11 as a disease gene and therapeutic target. The Biochemical journal 480, 1987–2008 (2023).

67. S. E. Compton et al., LKB1 controls inflammatory potential through CRTC2-dependent histone acetylation. Mol Cell 83, 1872–1886 e1875 (2023).

68. A. A. Widjaja et al., A Neutralizing IL-11 Antibody Improves Renal Function and Increases Lifespan in a Mouse Model of Alport Syndrome. J Am Soc Nephrol 33, 718–730 (2022).

69. D. Schumacher et al., A neutralizing IL-11 antibody reduces vessel hyperplasia in a mouse carotid artery wire injury model. Scientific reports 11, 20674 (2021).

70. S. A. Menzies et al., The sterol-responsive RNF145 E3 ubiquitin ligase mediates the degradation of HMG-CoA reductase together with gp78 and Hrd1. eLife 7, (2018).

71. A. Dolan et al., Genetic content of wild-type human cytomegalovirus. Journal of General Virology 85, 1301–1312 (2004).

72. R. J. Stanton et al., Reconstruction of the complete human cytomegalovirus genome in a BAC reveals RL13 to be a potent inhibitor of replication. Journal of Clinical Investigation 120, 3191–3208 (2010).

73. R. J. Stanton et al., Cytomegalovirus destruction of focal adhesions revealed in a high-throughput Western blot analysis of cellular protein expression. Journal of Virology 81, 7860–7872 (2007).

74. T. J. Blanchard, A. Alcami, P. Andrea, G. L. Smith, Modified vaccinia virus Ankara undergoes limited replication in human cells and lacks several immunomodulatory proteins: implications for use as a human vaccine. J Gen Virol 79 **( Pt** **5**), 1159–1167 (1998).

75. M. Law, G. L. Smith, Antibody neutralization of the extracellular enveloped form of vaccinia virus. Virology 280, 132–142 (2001).

76. W. W. Gierasch et al., Construction and characterization of bacterial artificial chromosomes containing HSV-1 strains 17 and KOS. J Virol Methods 135, 197–206 (2006).

77. B. Bushnell. vol. 2023.

78. A. Dobin et al., STAR: ultrafast universal RNA-seq aligner. *Bioinformatics (Oxford*, England) 29, 15–21 (2013).

79. S. Anders, P. T. Pyl, W. Huber, HTSeq--a Python framework to work with high-throughput sequencing data. *Bioinformatics (Oxford*, England) 31, 166–169 (2015).

80. M. D. Robinson, D. J. McCarthy, G. K. Smyth, edgeR: a Bioconductor package for differential expression analysis of digital gene expression data. Bioinformatics (Oxford, England) 26, 139–140 (2010).

81. G. C. McAlister et al., MultiNotch MS3 enables accurate, sensitive, and multiplexed detection of differential expression across cancer cell line proteomes. Anal Chem 86, 7150–7158 (2014).

82. W. Haas et al., Optimization and use of peptide mass measurement accuracy in shotgun proteomics. Mol Cell Proteomics 5, 1326–1337 (2006).

83. J. E. Elias, S. P. Gygi, Target-decoy search strategy for increased confidence in large-scale protein identifications by mass spectrometry. Nat Methods 4, 207–214 (2007).

84. J. E. Elias, S. P. Gygi, Target-decoy search strategy for mass spectrometry-based proteomics. Methods in Molecular Biology 604, 55–71 (2010).

85. R. Wu et al., Correct interpretation of comprehensive phosphorylation dynamics requires normalization by protein expression changes. Mol Cell Proteomics 10, M111 009654 (2011).

86. W. Kim et al., Systematic and quantitative assessment of the ubiquitin-modified proteome. Mol Cell 44, 325–340 (2011).

87. E. L. Huttlin et al., A tissue-specific atlas of mouse protein phosphorylation and expression. Cell 143, 1174–1189 (2010).

88. L. Kall, J. D. Canterbury, J. Weston, W. S. Noble, M. J. MacCoss, Semi-supervised learning for peptide identification from shotgun proteomics datasets. Nat Methods 4, 923–925 (2007).

89. W. Huang da, B. T. Sherman, R. A. Lempicki, Systematic and integrative analysis of large gene lists using DAVID bioinformatics resources. Nature protocols 4, 44–57 (2009).

90. D. R. Stirling et al., CellProfiler 4: improvements in speed, utility and usability. BMC Bioinformatics 22, 433 (2021).

91. J. A. Vizcaino et al., 2016 update of the PRIDE database and its related tools. Nucleic acids research 44, D447–456 (2016).

